# Directed Evolution of AAV Targeting Primate Retina by Intravitreal Injection Identifies R100, a Variant Demonstrating Robust Gene Delivery and Therapeutic Efficacy in Non-Human Primates

**DOI:** 10.1101/2021.06.24.449775

**Authors:** Melissa Kotterman, Ghezal Beliakoff, Roxanne Croze, Tandis Vazin, Christopher Schmitt, Paul Szymanski, Meredith Leong, Melissa Quezada, Jenny Holt, Katherine Barglow, Mohammad Hassanipour, David Schaffer, Peter Francis, David Kirn

## Abstract

Targeted AAV vectors are needed for safe and efficient delivery to and transduction of specific tissue target(s) in patients. Effective intravitreal delivery for retina gene therapy is not feasible with wildtype AAV. We employed directed evolution in nonhuman primates (NHP) to discover an AAV variant (R100) for intravitreal treatment of multiple target cells in the primate retina. R100 demonstrated superior transduction of human retinal cells compared to wildtype AAV. Furthermore, three R100-based gene therapeutics demonstrated safety, delivery, and durable pan-retinal expression of intracellular or secreted transgenes throughout the NHP retina following intravitreal administration. Finally, efficacy of R100-mediated delivery of therapeutic transgenes was demonstrated in patient-derived retinal cells (monogenic diseases) and in an NHP model of pathogenic retinal angiogenesis.

---

Recombinant adeno-associated viruses (AAV) are used as gene delivery vectors *in vivo* following insertion of a therapeutic transgene between the inverted terminal repeats (ITRs) in place of the viral *rep* and *cap* genes.^1^ Naturally-occurring (wildtype) AAV serotypes^2–8^ demonstrate transduction of cells *in vivo*, resulting in stable transgene expression for years in post-mitotic tissue.^9^ Furthermore, AAV-based gene therapeutics have yielded promising results in clinical trials.^10–20^ Recently, the U.S. Food and Drug Administration approved both Luxturna (voretigene neparvovec-rzyl), an AAV2-based gene therapy for the treatment of inherited blinding diseases caused by mutations in the *RPE65* gene, and Zolgensma (onasemnogene abeparvovec-xioi), an AAV9-based gene therapy for the treatment of spinal muscular atrophy.

However, several hurdles limit the clinical benefit of AAV in humans. First, wildtype vectors mediate inefficient delivery to most target tissues, necessitating very high doses and/or suboptimal delivery methods for patient treatment.^10–14, 20–22^ Second, for vector that reaches the intended tissue, target cell transduction and transgene expression can be highly inefficient.^23–25^ Third, dose-dependent inflammation and associated toxicities^15, 26^ leading to adverse events may occur with wildtype AAV due to the high doses required to achieve transduction in some tissues. For example, three patients who recently received a high intravenous dose of 3×10^14^ vg/kg of an AAV8-based gene therapy in a clinical trial for X-linked myotubular myopathy (NCT03199469) experienced progressive liver dysfunction that ultimately resulted in death. Fourth, pre-existing anti-capsid antibodies can limit efficacy pre-clinically and clinically.^27–30^

For treatment of the primate retina specifically, broad delivery to multiple cell types across the entire tissue has not been achievable with wildtype AAV capsids. Unfortunately, wildtype AAV capsids are not able to efficiently migrate through the inner limiting membrane (ILM) and superficial ganglion cell layer to achieve broad, efficient delivery to the posterior of the retina via an intravitreal injection.^31^ As a result, subretinal surgery has been the delivery method used most frequently in ocular AAV gene therapy clinical trials to date.^10, 11, 20, 32–36^ However, subretinal delivery efficiently transduces cells primarily in and closely surrounding the bleb region, which can reportedly account for less than 5% of the retina^14, 37^, and it results in retinal tears or detachments and other safety risks (including cataracts and thinning of the corneal stroma).^20, 35, 38^ Specifically, in clinical trials for Luxturna, 66% of subjects and 57% of eyes experienced ocular adverse reactions. The most common adverse events (defined as occurring in > 5% of subjects) were included conjunctival hyperemia, cataract, increased intraocular pressure, retinal tear, inflammation, macular hole, retinal deposit, irritation, pain, and maculopathy. ^20, 39^ In contrast, intravitreal injection is a safe, non-surgical procedure performed routinely in outpatient clinics for the delivery of protein therapeutics to treat common retinal diseases such as wet age-related macular degeneration (wet AMD). The most common adverse reaction following intravitreal injection is ocular inflammation, which typically resolves following treatment with corticosteroids.^40^ Importantly, the entire retinal surface can be treated by this method. Moreover, in contrast to subretinal surgery, the intravitreal delivery procedure does not result in anatomical detachment of the retina.

Directed evolution^41^ is a strategy to harness genetic diversification and selection processes to enable the creation and discovery of novel synthetic biologics. We applied directed evolution to generate novel AAV variants selected for the treatment of specific tissues in primates (Figure 1a). We hypothesized that applying this methodology to our diverse library of an estimated 100 million AAV *cap* gene variant sequences could lead to the selection of novel AAV variants capable of efficient delivery to specific tissues by clinically optimal routes. For retinal gene therapy, we used directed evolution for selection of vectors for administration by intravitreal delivery and subsequent transduction of the entire breadth of the retina (macula-to-periphery), including multiple target cell types such as photoreceptors, retinal pigment epithelial (RPE) cells, and ganglion cells. For identification of vectors that can be delivered by intravitreal administration to humans, the NHP eye is the most appropriate model due to similarities lacking in canine and rodent eyes. First, the NHP ILM more closely resembles the human thickness and composition, compared to dog and mouse ILM.^42, 43^ Also, the NHP eye and ocular structures are anatomically similar to the human eye, including the ocular contents, size and volume relationships, the presence of a macula, and the distribution and number of cone and rod photoreceptors. Given anatomical and physiologic similarities between NHP and human retinas and ILM, directed evolution was carried out in NHP. Previous attempts to develop next-generation AAV vectors have included evolution in mouse, dog, and NHP.^44–46^ To date, these attempts have demonstrated the potential applicability of directed evolution to address these challenges, but have yet to identify a variant capable of broad, efficient transduction of the primate retina following intravitreal administration. Intravitreal administration of AAV7m8, an AAV capsid evolved in mice, encoding a GFP transgene required a high dose of 5×10^12^ vg to achieve some transduction of the NHP retina, and RPE transduction was not observed.^47^ Proof-of-concept for NHP selection has previously been used to identify AAV variants capable of transduction of the retina, but as yet, these capsids have demonstrated neither widespread retinal transduction nor subsequent translatability to human therapeutics.^46^

**Figure 1.**
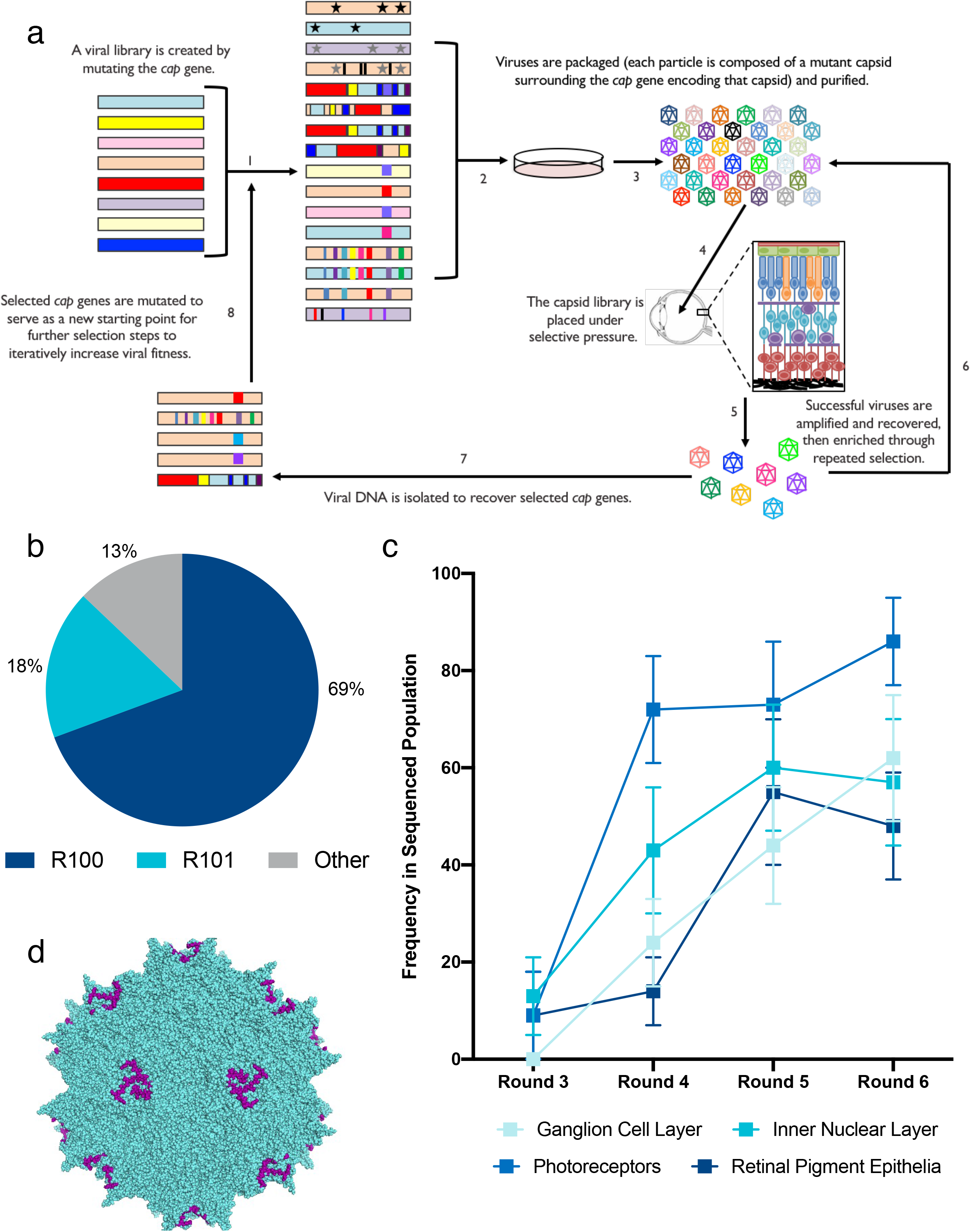
Directed evolution of AAV led to the discovery of a dominant motif capable of localization to all layers of the primate retina following intravitreal delivery. **a**, Directed evolution schematic. 1) plasmid library comprising 20+ combinations of DNA mutation techniques and *cap* genes is created; 2) viral library is packaged such that each particle comprises the mutant capsid surrounding the *cap* gene encoding that mutant capsid; 3) viral library is purified; 4) viral library is placed under selective pressure; 5) successful viruses are amplified and recovered; 6) successful viruses are enriched through repeated rounds of selection; 7) viral DNA is isolated to recover selected *cap* genes; 8) selected *cap* genes are mutated to serve as new starting library. **b**, Frequency of variant motifs within directed evolution Round 6 sequencing analysis. **c**, Frequency of the R100 variant motif found in separate retinal cell layers (ganglion cell layer, inner nuclear layer, photoreceptor layer, retinal pigment epithelial layer) in directed evolution Rounds 3-6. **d**, Representative three-dimensional model of R100. The AAV2-based variant contains an insertion of 10 amino acids (purple) at amino acid position 588. Error bars indicate the 95% confidence interval.

We identified a novel, evolved AAV capsid (R100) and subsequently carried out extensive characterization studies to confirm superior transduction efficiency of this capsid to the standard wildtype AAV used in the human retina (AAV2). Mechanistic analyses of R100 demonstrated that unique structural features of R100 lead to both improved ILM penetration, intraretinal distribution, and retinal cell transduction. In addition, vector characterization, safety, and efficacy studies were carried out in NHP and in normal and diseased human retinal cells. Finally, therapeutic constructs based on R100 were developed for both a rare monogenic recessive disease requiring intracellular protein expression and for large market diseases requiring secretion of a therapeutic protein.

## Results

### Directed evolution of AAV capsids in NHPs following intravitreal injection resulted in the creation and discovery of a synthetic AAV variant with a novel peptide insertion motif and point mutation

The directed evolution process was applied to discover AAV capsid variants capable of broadly transducing multiple layers of the primate retina following intravitreal (IVT) administration (Figure 1a). Briefly, a library of approximately 100 million unique synthetic variant AAV capsid sequences was created from 25 different sub-libraries using various molecular biology techniques and several different AAV serotypes as templates.^48–55^ The library was packaged in HEK293T cells to produce viral particles such that each virus particle was composed of a synthetic capsid shell surrounding the viral genome encoding that same capsid.^48, 49^ Variants within the library were then subjected to *in vivo* selective pressure techniques in NHP to mimic clinical intravitreal gene therapy treatment. That is, for each round of selection, the capsid library was injected into the vitreous humor. All synthetic libraries were injected for the first round of selection. After tissues were harvested from widespread regions of the retina, the genomes of capsids amplified from the tissue were then packaged as above for injection into another NHP for the next round of selection. This procedure was carried out for a total of six cycles with progressively lower vector titers. In order to increase diversity, additional mutagenesis techniques were applied to the remaining capsid sequences between rounds four and five.

Following capsid isolation for rounds 3-6 of the selection, sequencing was performed on individual clones within the library to determine the frequency of variants and sequence motifs within the capsid population. Five motifs were identified that each represented at least 5% of the sequenced population in multiple rounds, or at least 10% in a single round (Supplemental Figure S1). As the selection progressed, the motif within R100 increased to 69% of the capsid sequences remaining (Figure 1b), and this capsid also had the highest prevalence (40% - 86% in round 6) in all four retinal layers evaluated (Figure 1b-c). The R100 capsid contains a unique 10 amino acid peptide inserted at residue 588 and a point mutation in the VP1 region of the capsid (Figure 1d; Supplemental Figure S9).

### R100 efficiently transduces various human retinal cell types in vitro

To evaluate the ability of R100 to transduce human retinal cells, RPE cells, retinal ganglion cells (RGC), and photoreceptor culture models were generated and transduced with the R100 vector carrying an expression cassette containing a ubiquitously-active CAG promoter^56^ and the transgene for enhanced green fluorescent protein (*EGFP*) (R100.CAG-EGFP). Human RPE were generated from embryonic (hESCs) and induced pluripotent stem cells (iPSCs) (Supplemental Figure S2). To assess transduction of mature human RPE cells by R100.CAG-EGFP and the control AAV2.CAG-EGFP, both the hESC and iPSC-derived RPE cells were transduced at multiplicities of infection (MOI) of 50, 500, and 5,000 viral genomes per cell (vg/cell) and compared; AAV2 was selected as the control because it is the parental serotype of R100 and has clinical relevance in the retina.^10, 11, 13, 14, 20^ R100 was capable of significantly higher transduction efficiency and EGFP transgene expression than AAV2 in RPE cells from both sources (Figure 2a and 2b). By flow cytometric analyses, R100.CAG-EGFP transduced a statistically significantly higher percentage of target cells (∼65-75%) within the PMEL17+ population compared to AAV2.CAG-EGFP (∼20-25%) at an MOI of 5,000 (Figure 2a, b). Transgene expression kinetics revealed a more rapid onset of transgene expression by the R100 capsid compared to AAV2 (Figure 2c).

**Figure 2.**
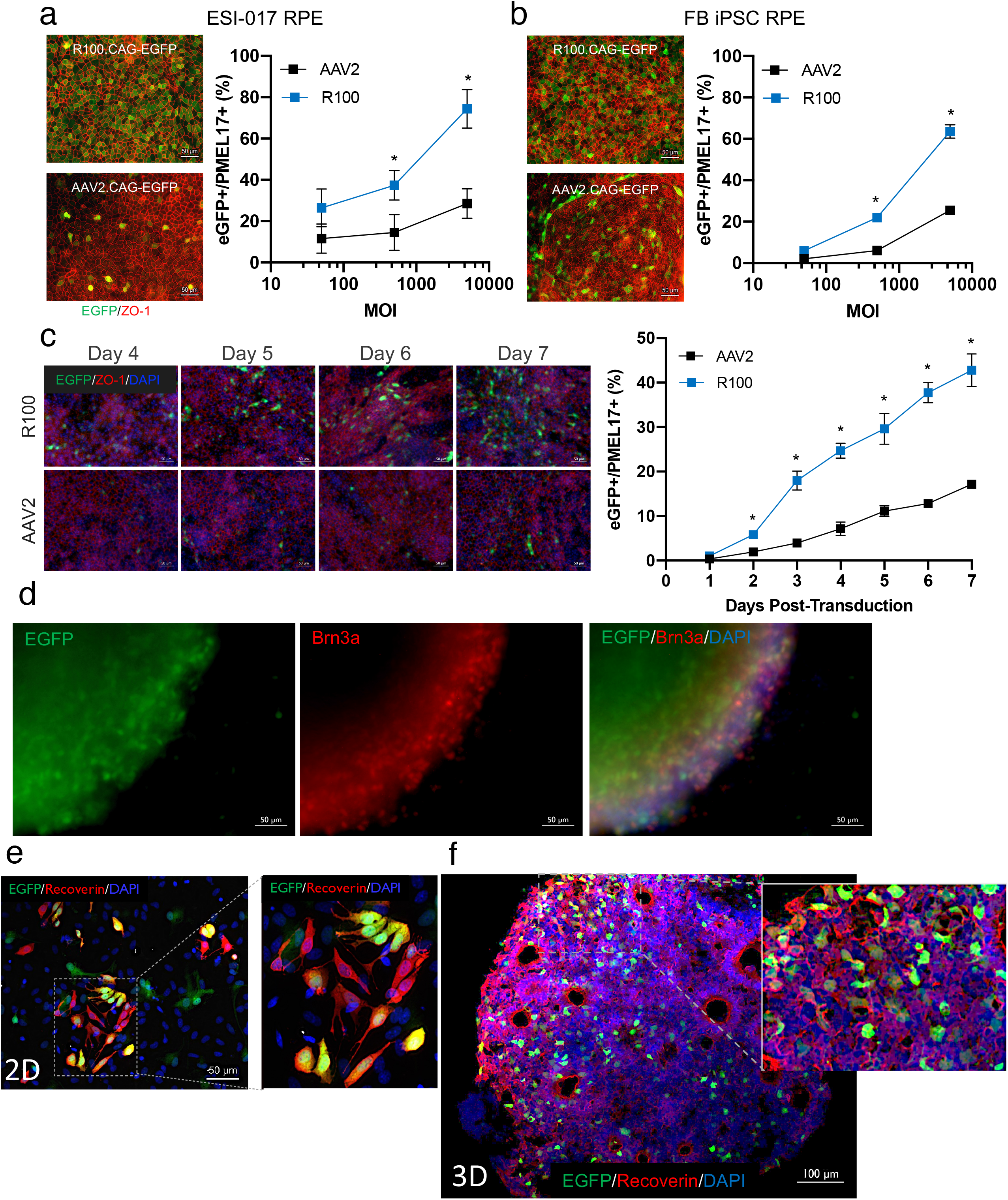
*In vitro* characterization of R100 in human cells. Representative images and quantitation of EGFP transgene expression (green) 6 days post-infection in **a**, hESC-derived and **b**, iPSC-derived RPE cells expressing ZO-1 (red) transduced with R100 or AAV2. Line graphs show EGFP+ cells in the PMEL17+ RPE cell population. **c**, EGFP expression kinetics following transduction by R100 or AAV2 in iPSC-derived RPE cells during the first 7 days post-infection. Representative images of EGFP transgene expression (green) following transduction with R100 in **d**, human Brn3a (red) expressing retinal ganglion cells 9 days post infection, **e,** adherent 2D human recoverin-expressing photoreceptors (red) 32 days post-infection and **f**, optic vesicles in suspension culture 14 days post-infection. **c**, **d**, **e**, **f**, Nuclei were counterstained with DAPI (blue). All images shown were cultures transduced at an MOI of 5,000, except for **c**, which were transduced at an MOI of 2,500. All quantitative measurements were carried out in n = 3; data presented as mean ± s.d.; * p < 0.05, compared to AAV2; two-tailed t-test.

Human RGCs were differentiated from normal fibroblast iPSCs according to published reports (Supplemental Figure S2).^57, 58^ To evaluate transduction of human RGCs by R100, retinal cultures were transduced with R100.CAG-EGFP at an MOI of 10,000. Cells were fixed 14 days post-transduction and examined for colocalization of Brn3a and EGFP by immunocytochemistry. EGFP expression was observed within the Brn3a positive population (Figure 2d), demonstrating the capacity of R100 to infect human RGCs *in vitro*.

2D and 3D human retinal cultures containing photoreceptors were generated from hESCs and iPSCs to evaluate transduction by R100 (Figure 2; Supplemental Figure S2). Both 2D and 3D retinal cultures, transduced at an MOI of 5,000 vg/cell, demonstrated EGFP expression colocalization with Recoverin positive photoreceptors (Figure 2e, 2f), thus indicating R100 transduction of human photoreceptors *in vitro*.

### Intravitreal administration of R100.CAG-EGFP in NHP resulted in widespread, robust transduction of multiple cell layers in both central and peripheral retinal regions

To evaluate transduction with R100 in the NHP retina following intravitreal administration, doses of 3×10^11^ vg/eye (n =3 NHP; Supplemental Table 2, Supplemental Figure S3) or 1×10^12^ vg/eye of R100.CAG-EGFP were administered to one or both eyes (n=9 NHP; Supplemental Table 2, Supplemental Figure S3). In-life fundus fluorescence, post-mortem immunofluorescence, and histological scoring were used to determine the distribution of transduction across the retina (macula, posterior pole, midperiphery and periphery) and the retinal cell types transduced (RPE, photoreceptors, and RGCs). No systemic or retinal toxicities were reported. Mild to moderate, transient corticosteroid-responsive anterior uveitis was observed in some animals, resolving within one to two weeks without sequelae. No posterior uveitis occurred in any animal between two weeks and six months.

Fundus fluorescence was utilized to monitor EGFP expression in-life, similar to previously published reports.^47, 59–61^ Overall, in-life analyses of EGFP expression following intravitreal injection of R100.CAG-EGFP showed robust, widespread transduction of the NHP retina as soon as three weeks after dosing in all retinal regions. In addition, expression was stable from eight weeks through six months post-administration. A dose response was observed, as animals receiving 1×10^12^ vg had more widespread and intense EGFP expression than those receiving the 3×10^11^ vg/eye dose (Supplemental Figure S3B; Supplemental Figure S5). Expression was observed in the fovea, macula, and periphery of the retina.

In-life fundus fluorescence images were confirmed and expanded upon through post-necropsy histology assessments. Fundus fluorescence enabled analysis of both the kinetics and distribution of transgene expression, though this methodology was not useful for quantitative assessments of total EGFP fluorescence due to camera autocalibration during image acquisition, and to diffraction of the excitation and emission spectra during travel through retinal tissue. Due to these limitations, extensive post-necropsy histology was also conducted. EGFP expression (visualized using direct EGFP fluorescence and an anti-GFP antibody) was observed broadly in both peripheral and central retina regions, including the fovea (Figure 3). RPE cells, photoreceptors, and RGCs were transduced throughout the retina (Figure 3b-d). In general, transgene expression was observed by fundus fluorescence in areas where high transgene expression was detected in at least two nuclei layers by immunofluorescence, which is a more sensitive method than in-life fundus fluorescence. Transgene expression from cone photoreceptors and RGCs approached 100% in assessed central foveal regions (Figure 3). Robust transduction efficiency, in many regions approaching 100%, was also noted in RPE cells in the periphery, midperiphery, and macula regions (Figure 3, Supplemental Figure S5), in contrast to reports of prior engineered vectors in mouse and NHP.^46, 47^ Likewise, in the peripheral retina, widespread transduction of RPE cells, photoreceptors, and RGCs was also observed (Figure 3, Supplemental Figure S5).

**Figure 3.**
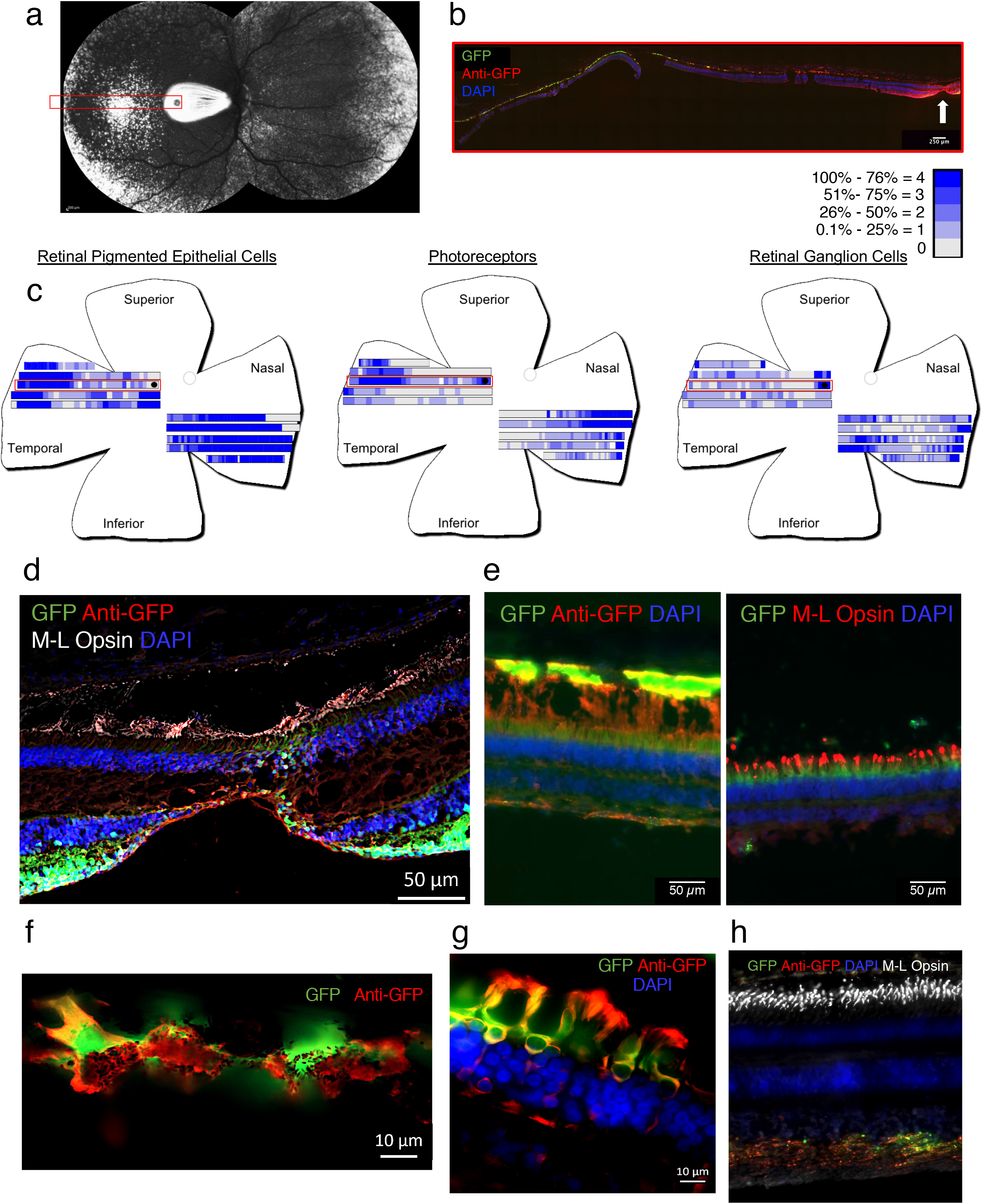
Comparison of fundus fluorescence, IHC imaging, and IHC quantitation demonstrates consistent and robust R100-mediated transgene expression following intravitreal delivery. NHPs received a dose of 1×10^12^ vg/eye of R100.CAG-EGFP. **a**, Fundus fluorescence imaging 3 weeks post-administration showed EGFP expression in the central, mid-periphery, and periphery of the retina (white). **b**, Representative image of a section of the retina (red rectangle in **a)** where robust EGFP expression is seen in multiple layers of the retina. The white arrow indicates location of the fovea. **c**, IHC scoring visualization. EGFP expression in the RPE layer (left), PR layer (middle), and RGC layer (right) was quantified for each visual field assessed and depicted as shades of blue corresponding to the percentage of EGFP+ cells within each retinal layer. The red rectangles represent the corresponding region in **a** and **b**. **d**, Representative image of transduced cells in the central retina. 20x image where robust EGFP expression is present in both cone PR (white) and RGC in the fovea. **e**, Representative images of robust EGFP expression in peripheral photoreceptors (left) and co-localized with opsin (right) **f**, High magnification representative image of robust EGFP expression present in RPE. **g**, High magnification representative image of robust EGFP expression present in the outer segments of PR. **h**, Representative image of robust GFP expression present in the axons of RGCs. EGFP expression (green) is also detected by an anti-GFP antibody (red) in all images. Nuclei were counterstained with DAPI (blue) except in **f**. Scale bar sizes are noted on each image.

Despite the known immunogenicity of EGFP, long-term in-life evaluations were undertaken to determine the kinetics of R100-mediated EGFP expression *in vivo*. NHPs were evaluated with in-life fundus fluorescence at various timepoints through six months post-administration (Figure 4, Supplemental Table 2, Supplemental Figure S3, S4). Detectable EGFP expression was observed as early as one-week post-administration of R100.CAG-EGFP (Figure 4c, Supplemental Figure S4), and expression significantly increased between one and three weeks post-administration (Figure 4c-d, Supplemental Figure S4). Durable and stable transgene expression was confirmed through six months post-administration (Figure 4e-g, Supplemental Figure S3, S4).

**Figure 4.**
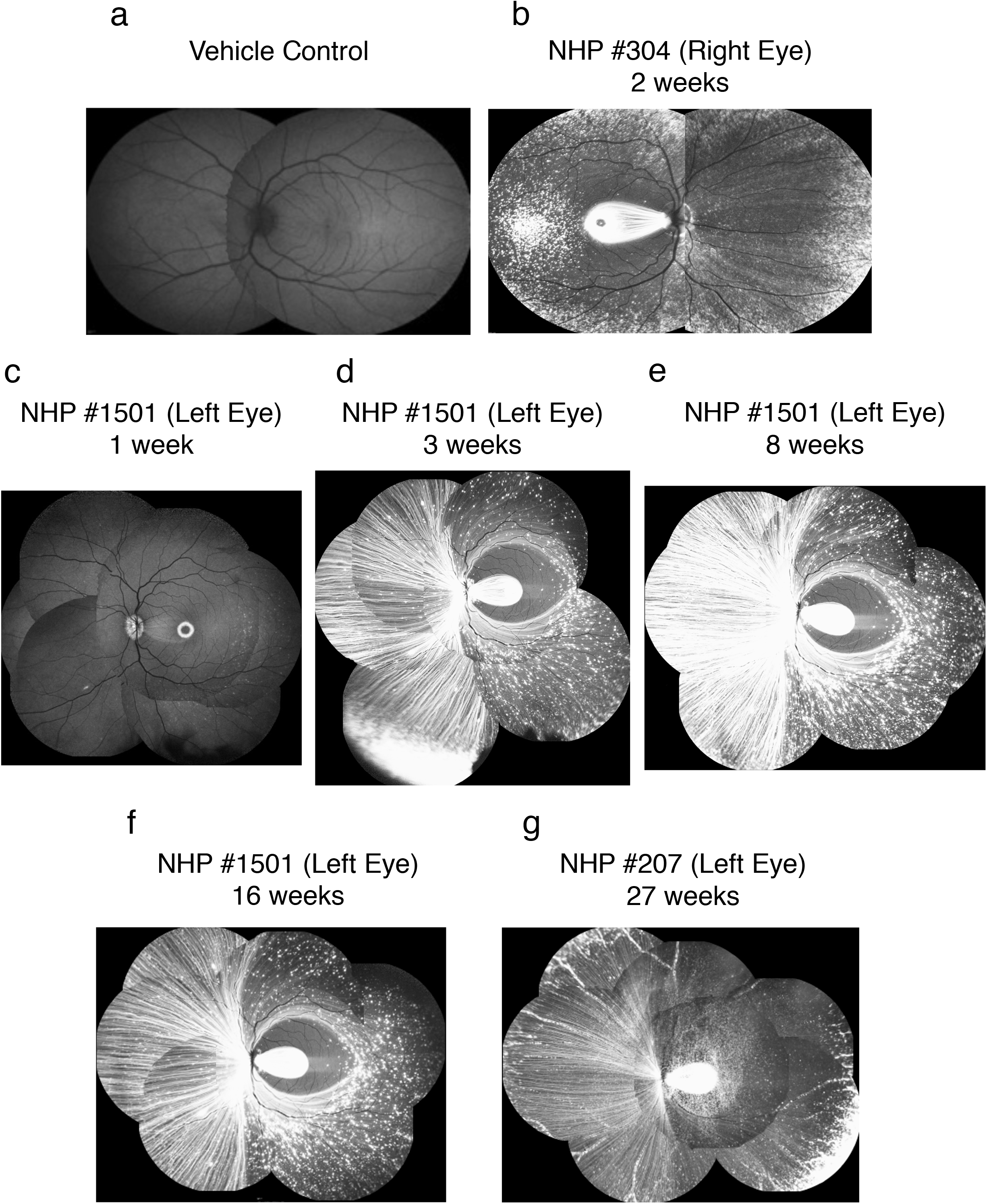
Representative fundus fluorescence images demonstrate robust R100-mediated transgene expression following intravitreal delivery to multiple NHP eyes. NHPs received a dose of 1×10^12^ vg/eye of R100.CAG-EGFP. EGFP expression (white) is present as early as 1 week post-administration and increases in intensity and distribution over time. **a**, vehicle control. **b**, NHP ID 304 right eye, 2 weeks post-administration. **c**, NHP ID 1501 left eye, 1 week post-administration. **d**, NHP ID 1501 left eye, 3 weeks post-administration. **e**, NHP ID 1501 left eye, 8 weeks post-administration. **f**, NHP ID 1501 left eye, 16 weeks post-administration. **g**, NHP ID 207 left eye, 27 weeks post-administration.

In summary, intravitreal injection of R100.CAG-EGFP showed robust, widespread transduction of the NHP retina, and transgene expression was observed at each time point assessed by fundus fluorescence throughout the duration of the study in all animals evaluated (Figure 3, Figure 4, Supplemental Figure S3). It is noted that cell-to-cell transfer of proteins, including GFP, between retinal cells has previously been reported.^62^ Although cellular localization of vector genomes and transcripts was not within the scope of this study, it is unlikely that cell-to-cell transfer would account for the widespread distribution of GFP described here. In addition, the breadth and intensity of EGFP expression observed by fundus fluorescence and immunofluorescence imaging using R100 by intravitreal injection was far superior to the transduction reported for any other wild-type serotype or variant of AAV reported to date.^47, 59, 60^

### R100 capsid peptide insertion results in reduced binding to a major component of the ILM, increased utilization of a broadly expressed glycan, and enhanced RPE cell transduction

We explored the mechanism(s) involved in the enhanced passage of R100 through the ILM barrier, plus enhanced transduction of human RPE cells, through *in silico* modeling and experimentation. The peptide insertion in the R100 capsid does not bear homology to any known mammalian peptides. Therefore, molecular modeling of the R100 peptide insertion and surrounding amino acids was performed to predict the structural implications of the peptide insertion. Within the wild-type AAV2 capsid, amino acids R585 and R588 lie in close proximity (Figure 5a), and are precisely positioned to interact with the heparan sulfate proteoglycan.^63, 64^ The R100 peptide insertion is predicted to have an extended helical structure, and the length and helical tendency of this sequence sterically precludes the close proximity of R585 and R588 required for heparin binding (Figure 5b). The presence of additional amino acids between positions 587 and 588 of the AAV2 capsid sequence has been shown previously to disrupt binding to heparan sulfate,^47, 53, 65^ therefore reducing affinity. The reduction in affinity for heparan sulfate was confirmed through binding to and elution from a heparin affinity column (Figure 5c). Heparan sulfate is present within the primate ILM,^31^ and therefore R100 has reduced binding affinity to a critical ILM component.

**Figure 5.**
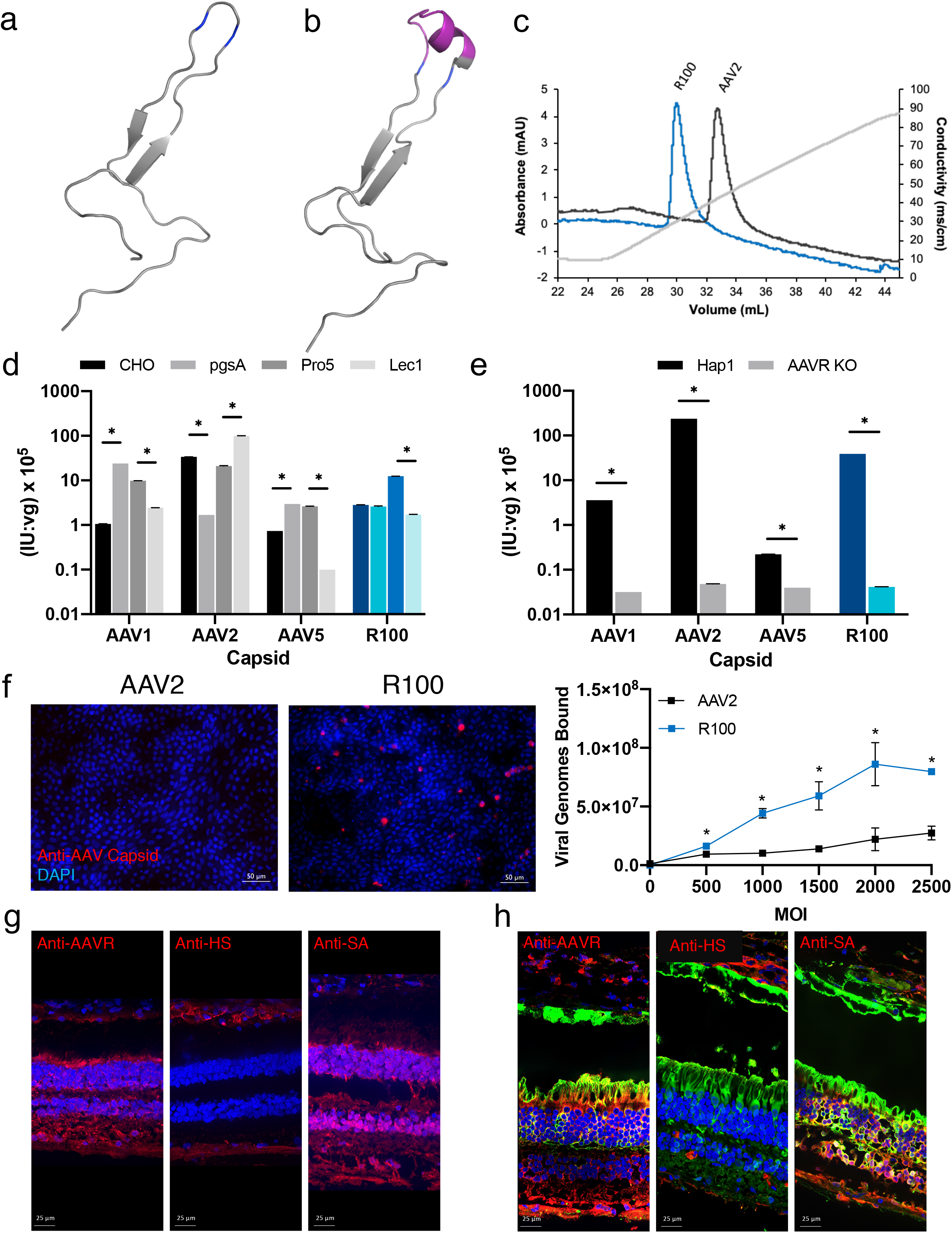
Mechanistic Analysis of R100 Transduction. Molecular models of **a**, AAV2 and **b**, the R100 peptide sequence (magenta) constrained by the location of amino acids R585 and R588 (blue) within the intact capsid. **c**, AAV2 and R100 vectors bound to and eluted from a heparin affinity column. R100 capsid elutes at a lower conductivity compared to AAV2, demonstrating a decrease in heparin affinity. **d**, AAV vectors transduced a panel of cell lines: CHO, pgsA (lacking surface glycosaminoglycans), Pro5, and Lec1 (lacking sialic acid). R100 demonstrates a shift in cell surface receptor dependence from heparan sulfate proteoglycans to sialic acid. **e**, AAV vectors were used to transduce Hap1 and AAVR KO Hap1 cell lines. All vectors demonstrate a cell surface receptor dependence for AAVR. **f**, Representative images of AAV2 and R100 binding to human iPSC-derived RPE, visualized using an anti-AAV2 capsid antibody and quantification of the number of viral genomes bound to human iPSC-derived RPE. Representative images of **g**, cell layer localization of AAVR, heparan sulfate, and sialic acid and **h**, colocalization of R100-mediated EGFP expression and cell surface receptors within the NHP retina. Error bars indicate standard deviation (n = 3); * p < 0.05, pgsA compared to CHO (d); Lec1 compared to Pro5 (d); AAVR compared to Hap1 (e); R100 compared to AAV2 (f); two-tailed t-test. Cell surface receptor expression is detected by an antibody (red) in all images. Nuclei were counterstained with DAPI (blue). Scale bar sizes are noted on each image. HS = heparan sulfate; SA = sialic acid.

The dependency of R100 on known AAV glycan co-receptors was then evaluated to better understand the mechanism(s) of cell entry. R100 and natural AAV serotypes AAV1, AAV2, and AAV5 were administered to several CHO-K1 cell lines, including pgsA cells (a CHO variant deficient in xylosyltransferase, leading to a lack of surface glycosaminoglycans),^66^ Pro5 cells (a CHO derivative proline auxotroph, parental line for Lec1),^66^ and Lec1 cells (a Pro5 variant deficient in GlcNAc glycosyl transferase, leading to a lack of surface N-glycans).^67^ The receptor dependencies of natural AAV serotypes AAV1 (α-2,3 and α-2,6 N-linked sialic acids),^68^ AAV5 (α-2,3 N-linked sialic acid),^69^ and AAV2 (heparan sulfate proteoglycans)^70^ were confirmed (Figure 5d). AAV2 is the parental serotype of R100, so it was anticipated that R100 would use the same primary receptor as AAV2. However, R100 dependency on heparan sulfate was reduced (although not completely ablated) compared to AAV2. In addition, R100 may require one or more of the deficient N-glycans (and potentially other as yet unidentified, ligands) to achieve transduction, an unanticipated result. R100 did, however, maintain dependence on the AAV receptor (AAVR),^71^ as demonstrated by a decrease in transduction of Hap1 cells lacking AAVR compared to normal Hap1 cells (Figure 5e).

The reduction of R100 affinity to heparan sulfate proteoglycans and gain of function affinity for an alternative glycan as a receptor have biological relevance in the retina. Improved transduction of human RPE cells (Figure 2) correlates with the enhanced binding of R100 to human RPE cells compared to AAV2 (Figure 5f). The enhanced binding of R100 is likely due to increased utilization of an alternative glycan receptor that is in abundance in the retina. In the NHP retina, heparan sulfate is present only in the ILM and the RGC layer (Figure 5g). However, sialic acid, for example, is ubiquitously present in all target cell layers, including photoreceptors and RPE cells (Figure 5g). R100-mediated EGFP expression co-localizes with AAVR and sialic acid within the NHP retina (Figure 5h), furthering a potential correlation between these receptors and R100 transduction. Of note, utilization of an additional glycan such as sialic acid alone is likely not sufficient for broad transduction of the retina, as evidenced by the lack of transduction by AAV1 and AAV5 following intravitreal administration when the ILM is intact.^31^ Therefore, the combination of low-affinity heparan sulfate binding, enhanced additional glycan binding, and the specific sequence of the peptide insertion in R100 all appear to be involved with the enhanced *in vivo* transduction following intravitreal injection.

### R100-mediated delivery and expression of Rep1 transgene results in functional and morphological correction of human retinal pigment epithelial cells derived from choroideremia patients

To explore efficacy of R100-based gene therapy to treat a monogenic recessive disease, an *in vitro* disease model using iPSC-derived RPE cells from choroideremia (CHM) patients was generated. Two CHM patient fibroblast samples were reprogrammed to iPSCs, then differentiated into functional, mature RPE cells (Supplemental Figure S6). Lack of Rep1 protein in CHM patients has been shown to correlate with cellular defects in Rab27a trafficking and prenylation, which subsequently leads to progressive degeneration of vision.^72–74^ Two weeks after CHM RPE cells were transduced with R100.CAG-cohREP1 (4D-110), Rab27a trafficking from cytoplasmic regions to target membranes was normalized (Figure 6a, 6c), correlating with restitution of the normal cellular RPE phenotype (Figure 6b).

**Figure 6.**
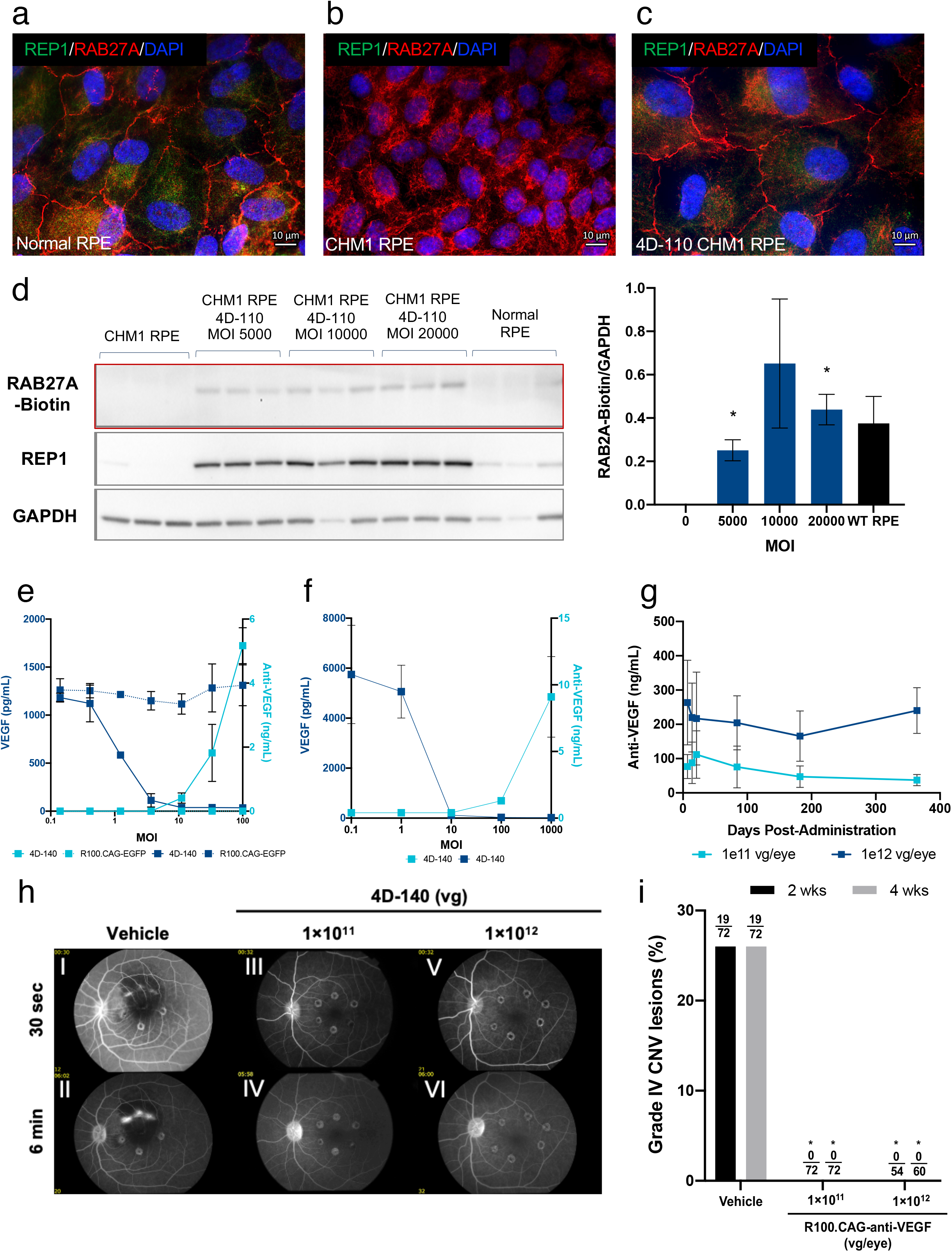
Expression and activity of R100 expressing therapeutic transgenes. **a**, normal iPSC-derived RPE cells **b**, CHM1 RPE, and **c**, CHM1 RPEs transduced by 4D-110 at an MOI of 5,000, evaluated for the presence of REP1 (green), and RAB27A (red) proteins. Nuclei counterstained with DAPI (blue). **d**, Level of REP1 protein by western blot analysis and incorporation of a biotinylated prenyl donor as a measure of prenylation in cell lysates from untransduced cells and 4D-110 transduced cells. Graph illustrates the average band intensity of biotinylated RAB27A relative to GAPDH for each sample. All quantitative measurements were n = 3; data presented as mean ± s.d.; * p < 0.05, compared to untransduced CHM RPE; two-tailed t-test. **e**, normal human RPE and **f**, RGCs transduced by 4D-140 assayed for VEGF or anti-VEGF by ELISA. Line graph illustrates a dose-dependent decrease in the level of free VEGF and a concomitant increase in the level of free anti-VEGF protein. Error bars indicate the standard deviation of **e**, duplicate transduction wells or **f**, triplicate transduction wells. Laser-induced model of CNV in primates. NHP (n=6) were dosed bilaterally IVT with vehicle or 4D-140 and CNV induced 42 days post-dose. **g**, levels of anti-VEGF scFv were assessed in aqueous humor at various time points. **h**, representative images two weeks post-laser of early and late phase fluorescein angiograms of eyes treated with vehicle (I, II), 1×10^11^ vg/eye (III, VI), and 1×10^12^ vg/eye (V, VI) of 4D-140. **i**, incidence of grade IV lesions at two (black) and four (gray) weeks post-laser treatment. Numbers indicate total grade IV lesions above total assessable lesions per group. * p < 0.0001 compared to vehicle using Fisher’s exact test.

A functional assay was developed to assess the ability of R100-mediated REP1 protein expression to restore normal Rab27a GTPase prenylation as determined by Western blot. Treatment of CHM RPE cells with 4D-110 resulted in both Rep1 protein and prenylated Rab27a levels at or above that of normal RPE (Figure 6d), representing functional correction of the prenylation defect in choroideremia patient-derived RPE cells.

### R100-mediated delivery and expression of anti-VEGF transgene results in functional activity in human retinal cells in vitro and pathology correction in an NHP model of choroidal neovascularization

To explore efficacy of R100-based gene therapy with a secreted transgene product for treatment of complex retinal diseases such as wet AMD, a therapeutic payload was designed and engineered to express an anti-VEGF single chain variable fragment (scFv). Normal human iPSC derived RPE and RGC cultures were transduced with R100.CAG-antiVEGF scFv (4D-140) at MOIs ranging from 0.15 to 100; R100.CAG-GFP was used as a negative control. Media samples were collected from four to seven days post-transduction and assayed for free (unbound) VEGF and free anti-VEGF scFv by ELISA. RPE cells and RGCs endogenously produce VEGF, and in both cell types treatment with 4D-140 resulted in decreased concentrations of free secreted VEGF and increased concentrations of free secreted anti-VEGF scFv in a dose-dependent manner (Figure 6e and f). At MOIs of 10 and above, VEGF was largely sequestered in treated RGC cultures. These results show that the 4D-140 product (and thus the R100 capsid) leads to expression and secretion of functional anti-VEGF scFv in *in vitro* models of both human RPE cells and RGCs.

To assess expression and efficacy *in vivo*, NHPs were dosed IVT with either 1×10^11^ or 1×10^12^ vg/eye of 4D-140 or vehicle control, and anti-VEGF scFv was assessed over time in aqueous humor by ELISA. Anti-VEGF scFv was detected in aqueous humor of all treated animals in a dose-dependent manner (Figure 6g). Expression was detected as early as 7 days post-dosing with levels maintained through 12 months post-dosing (Figure 6g). R100-mediated expression was observed in all NHP eyes at all time points for both doses evaluated. Six weeks post-administration, NHPs underwent laser photocoagulation to the perimacular region of the retina to induce choroidal neovascular lesions. Two and four weeks after administration of the laser, lesions were assessed by fluorescein angiography, and the number of total assessable lesions per eye and the number of grade IV CNV lesions (defined by hyperfluorescence early or mid-transit with late leakage extending beyond borders of the treated area) per eye was determined. Representative images of the incidence of grade IV lesions in vehicle control-treated animals was 26% (19/72) at both timepoints, which is within the previously published range of performance for this model (25% - 40%).^75–77^ In contrast, no grade IV lesions were detected in any animals treated with 4D-140 at either 1×10^11^ or 1×10^12^ vg/eye (0/72 and 0/54 at two weeks; 0/72 and 0/60 at four weeks, respectively; Figure 6i), demonstrating a significant improvement compared to vehicle controls. No significant product-related toxicities were observed with 4D-140 at either dose level, as determined by clinical observations (Supplementary Figure S7). Finally, bilateral administration of 4D-140 was well-tolerated, with minor or mild inflammation in some eyes that received the 1×10^12^ vg/eye dose (Supplementary Figure S7).

## Discussion

The data described here demonstrate that the synthetic R100 AAV capsid variant offers promise for intravitreal gene therapy of both monogenic and complex diseases of the retina, including for the expression of both intracellular and secreted protein therapeutics. Efficacy was demonstrated both in human patient-derived retinal cell models *in vitro* and in a primate model of complex angiogenic diseases *in vivo*. This novel capsid variant resulted in robust transduction of human RPE cells, RGCs, and photoreceptors *in vitro* and primate (NHP) RPE, photoreceptors, and RGCs across central, mid-peripheral, and peripheral regions of the retina *in vivo* following intravitreal administration. Safety, transgene expression, and low cell-mediated immunogenicity were demonstrated across three GLP toxicology and biodistribution studies in NHP. R100 was identified through a directed evolution screen to select capsid variants that were able to traverse the ILM and transduce multiple retinal cell types broadly across the entire surface area of the retina following intravitreal administration in primates. The changes to the R100 capsid resulted in a gain of function to utilize additional cell surface glycans for binding and entry into cells, an unanticipated mechanism that has not previously been reported for AAV2-based variants, in addition to reduced binding to the heparin sulfate present in the ILM. Superiority of R100 to AAV2 wildtype capsid vector transduction was demonstrated in human cells *in vitro*. To our knowledge, this is the first demonstration of directed evolution and full characterization of a gene delivery vehicle in non-human primates and in human target cells confirming the desired delivery, transduction, efficacy and safety phenotype suitable for human therapy. This finding may have implications for AAV capsid discovery for the treatment of other tissues and with other routes of administration in primates for use in humans, as well as for other viral vectors. For example, these results may be predictive for successful selections for novel capsids for intravenous low dose delivery to heart and skeletal muscle, and for aerosol delivery to the lung airway epithelium.^78, 79^

Previous publications that have described directed evolution of the AAV capsid in murine, canine, and NHP eyes *in vivo* demonstrate proof-of-concept that *in vivo* selections can generate novel capsids.^25, 45, 47, 80^ These previous selections were performed with limited capsid library diversity (both total diversity and the types of mutations included) compared to this study.^25, 45, 47, 80^ To date, none of the previously identified capsid variants have demonstrated broad, robust, and durable expression across multiple cell types and cell layers within the primate retina, or beyond the fovea. The mouse evolved AAV7m8 vector required high doses to achieve transduction in NHP, was not capable of transducing RPE, and did not robustly transduce outside of the fovea.^47^ This is further evidenced by the use of a 2×10^12^ vg/eye dose to show efficacy in the NHP model of laser induced choroidal neovascularization.^75^ Current Phase I clinical data with this capsid has demonstrated potential efficacy in patients, but has required administration in some cases of long term immunosuppressives to control ocular inflammation (NCT03748784). In contrast, R100 mediated robust EGFP expression throughout the primate retina and in all cell layers (including RPE) at a dose of 1×10^12^ vg/eye or lower, and R100 mediated anti-VEGF scFv expression resulted in complete suppression of grade IV lesions at a lower 1×10^11^ vg/eye dose in the NHP model of laser induced choroidal neovascularization. The R100 capsid efficiently delivered intracellular and secreted therapeutic transgenes and resulted in phenotypic correction of a monogenic retinal disorder in a human RPE cell model and a complex retinal disorder in an NHP disease model. We therefore believe that directed evolution, vector characterization, and vector safety and efficacy studies should be carried out in human cells *in vitro* and in NHP *in vivo* for better human clinical translation.

In contrast to these previous proof-of-concept demonstrations of directed evolution feasibility, this study reports efficacy with the evolved, synthetic vector R100 for a rare disease in patient-derived retina cells *in vitro* and using a secreted anti-angiogenic factor both *in vitro* and in the primate model of pathologic angiogenesis. The significant reduction in incidence of grade IV CNV lesions with doses as low as 1×10^11^ vg/eye demonstrates the robust ability of R100 platform to deliver effective therapeutics transgenes following IVT administration in primates. Significantly superior transduction efficiency to wildtype AAV2 was demonstrated *in vitro* in human retina target cells, as well as superior transduction to what has been described with AAV2 in NHP, *in vivo*. In summary, this publication reports on a robust primate vector directed evolution discovery program and a proposed mechanism for the increased efficacy, and it describes full characterization of a novel evolved vector and associated gene therapy products in both normal and diseased primate and human model systems.

These data also demonstrate that directed evolution can simultaneously improve multiple phenotypic characteristics of a viral vector. With R100, we demonstrated improvements over parental AAV2 including its delivery to target tissues by an optimal route of administration (intravitreal), its ability to penetrate through a physical barrier to AAV (the ILM), and its ability to transduce and express transgenes from a target human or primate cell type (RPE and PR cells). We report that the presence of the peptide insertion in the R100 capsid simultaneously decreased affinity for heparan sulfate, presumably allowing for penetration through the heparan sulfate-rich ILM barrier, while increasing affinity for target retinal cells through enhanced binding to additional surface glycan receptors. Of note, this peptide insertion is repeated 60 times across the R100 capsid surface.

Two products including the R100 capsid have recently been translated into clinical trials for inherited retinal diseases that primarily involve different retinal cell layers: choroideremia (*Rep-1* delivery to RPE cells; NCT04483440) and X-linked retinitis pigmentosa (*RPGRorf15* delivery to PR cells; NCT04517149). These two therapeutics have been evaluated by unilateral and bilateral intravitreal administration to NHPs in three separate GLP toxicology and biodistribution studies, and the results supported the filing of Investigational New Drug applications and clinical trial testing. Human clinical trials are underway to determine the safety, pharmacodynamics, and efficacy (including through serial visual field testing and optical coherence tomography scans) of these R100 vector-based therapeutics. The consistency in the biodistribution, gene expression, vector clearance, and minimal immune response between independent products and across preclinical studies supports the development of additional products using R100 as a platform vector to treat diverse retinal disorders. R100 was developed to efficiently transduce multiple cell types across all layers of the retina. This quality allows for the use of a single capsid for the treatment of diseases of the RPE, photoreceptors, subsets of photoreceptors, or RGCs, through delivery of transgenes encoding intracellular therapeutic proteins, transgenes encoding secreted therapeutic proteins, homology-directed repair payloads, silencing/knock-down payloads, gene editing nucleases, or transgenes encoding optogenetic channel proteins. If expression of a particular therapeutic transgene needs to be restricted to a specific cell type or cell types, the use of cell-specific promoters and/or other regulatory elements can be incorporated into the therapeutic payload. This concept enables a streamlined capsid preclinical development program and more rapid enablement of multiple therapeutic products for both monogenic and complex disorders.^78, 79, 81, 82^

We describe here that directed evolution of AAV in NHP can be used not only to discover novel variants, but also a novel capsid variant capable of transduction of relevant target cells in NHP by the preferred clinical route, with a favorable safety profile, and that efficient transgene expression and efficacy can be achieved in human target cells as well. These principles should be replicable for vectors targeting other cells or tissue types by optimal routes of administration in NHP and humans.^78, 79, 81, 82^ These data demonstrated the feasibility of evolving vectors with improved trafficking through bodily fluid(s), traversing of physical barriers, migration through dense tissues, and transduction of multiple cell types. These principles can be applied to other neural and sensory tissues of similar structure and/or cell types, such as the brain, spinal cord, and ear. In addition, directed evolution can be used to discover vectors that are resistant to pre-existing antibodies in the human population.^49^ Finally, we demonstrated that directed evolution can be used to simultaneously optimize several phenotypic features and multiple steps in the pathway from injection to therapeutic efficacy. This report thus represents the first example of the application of directed evolution to develop an improved AAV vector specifically for better human gene therapies through demonstration of utility, superiority to AAV2, and mechanism of transduction.

## Online Methods

### Cell Lines and Plasmids

Cell lines were cultured at 37°C and 5% CO_2_ and, unless otherwise mentioned, were obtained from the American Type Culture Collection. HEK293T, HEK293 (National Research Council of Canada), and 2v6.11 (Johns Hopkins University) cells were cultured in Dulbecco’s modified Eagle’s medium (DMEM) supplemented with 10% fetal bovine serum (Hyclone; GE Healthcare Life Sciences) and 1% penicillin/streptomycin (ThermoFisher). The human embryonic stem cell line ESI-017 (ESI BIO), a human fibroblast-derived induced pluripotent stem cell line, FB-iPSC, (System Biosciences), and CHM-iPSC lines were cultured on Vitronectin XF (Stem Cell Technologies) in mTeSR-1 maintenance medium (Stem Cell Technologies). Human pluripotent stem cell (hPSC) cultures were sub-cultured using Gentle Cell Dissociation Reagent (Stem Cell Technologies), every four to five days at 70-80% confluence. Human primary umbilical vein endothelial cells (HUVEC) (Lonza) were cultured in EBM-2 media (Lonza). ARPE19 cells were maintained in DMEM:Nutrient Mixture F-12 and sodium pyruvate (ThermoFisher), supplemented with GlutaMAX, and 15 mM HEPES (ThermoFisher) and 10% fetal bovine serum (GE). Human hepatocytic carcinoma HepG2 cells were cultivated in Eagle’s Minimum Essential Medium, supplemented with 10% fetal bovine serum (GE). CHO K1 and CHO pgsA cells were cultured in F-12K medium (ATCC) supplemented with 10% fetal bovine serum (GE) and 1% penicillin/streptomycin (Invitrogen). Pro5 and Lec1 cells were cultured in MEM-alpha medium (GE) supplemented with 10% fetal bovine serum (GE) and 1% penicillin/streptomycin (Invitrogen). Hap1 and Hap1 KO cells were cultured in Iscove’s modified Dulbecco’s medium (Gibco) supplemented with 10% fetal bovine serum (GE) and 1% penicillin/streptomycin (ThermoFisher).

For generation of the CAG-EGFP plasmid, a DNA sequence totaling 3,029 bp containing the AAV2 5’ inverted terminal repeat (ITR), CAG promoter, enhanced GFP transgene, polyadenylation sequence, and AAV2 3’ ITR was synthesized *de novo* by GenScript. The target sequence was synthesized as oligo fragments, then assembled to generate the full target sequence. The full target sequence was ligated into the backbone plasmid containing an ampicillin resistance gene using BamHI and HindIII restriction sites. The full target sequence was confirmed using Sanger sequencing.

For generation of the CAG-cohREP1 plasmid, a DNA sequence totaling 4,246 bp containing the AAV2 5’ inverted terminal repeat (ITR), CAG promoter, codon optimized Rep1 cDNA transgene, polyadenylation sequence, and AAV2 3’ ITR was synthesized *de novo* by GenScript. The Rep1 cDNA transgene was codon optimized using GenScript’s proprietary OptimumGene codon optimization algorithm. The target sequence was synthesized as oligo fragments, then assembled to generate the full target sequence. The full target sequence was ligated into the backbone plasmid containing an ampicillin resistance gene using BamHI and HindIII restriction sites. The full target sequence was confirmed using Sanger sequencing.

For generation of the CAG-Anti-VEGF scFv plasmid, a DNA sequence totaling 3,142 bp containing the AAV2 5’ inverted terminal repeat (ITR), CAG promoter, codon optimized anti-VEGF scFv cDNA transgene preceded by a human Ig kappa light chain signal peptide (synthesized *de novo* by GeneArt/ThermoFisher), polyadenylation sequence, and AAV2 3’ ITR was constructed by ligation of the anti-VEGF scFv sequence into a backbone plasmid containing a kanamycin resistance gene using NheI and MluI restriction sites. The full target sequence was confirmed using next generation sequencing.

### Library Generation and Viral Library Production

Plasmids containing *cap* gene libraries consisting of random mutagenesis of the AAV2 *cap* gene, the AAV2 *cap* gene containing random peptide insertions, and the AAV2 *cap* gene containing randomized hypervariable loop regions, and shuffled DNA from the wild-type AAV1, AAV2, AAV4, AAV5, AAV6, AAV8, AAV9 cap genes were generated as previously described.^49, 51, 53, 54^ Plasmids containing *cap* gene libraries consisting of random mutagenesis of the *cap* genes of other AAV serotypes and random peptide insertions/substitutions into the *cap* genes of other AAV serotypes were generated using similar methods to those previously described.^49, 53^ Plasmid libraries were sequenced using Sanger sequencing to determine the percentage of unique sequences and the percentage of functional variants (lacking stop codons and/or frame shift mutations before and after viral packaging.

Libraries were packaged via triple transfection of HEK293T cells using polyethylenimine (PEI; Polysciences, Inc.) at plasmid quantities and ratios designed such that the majority of cells receive a single member of the library.^48^ The libraries were purified by iodixanol gradient centrifugation and Amicon filtration, as previously described.^49, 51, 53, 54^ DNase-resistant genomic titers were determined via quantitative PCR.^49, 51, 53, 54^ For the initial round of selection, libraries were combined in equal viral genome quantities. Following completion of directed evolution, the *cap* gene sequence for R100 was inserted into the pXX2 recombinant AAV packaging plasmid using NotI and HindIII, as previously described.^48^

### Animal Use, Care, and Handling

For directed evolution and R100 Vector Characterization studies, drug-naive, neutralizing antibody (NAb) negative cynomolgus macaques (*Macaca fascicularis*) of both sexes in the age range of 2-14 years old were obtained. For the CNV study, drug-naive, NAb negative African green monkeys (*Chlorocebus sabaeus*) of both sexes in the age range of 4-13 years old were humanely procured from the healthy wild population. All animals were used in accordance with ARVO Statement for the Use of Animals in Ophthalmic and Vision Research. All aspects of the animal studies were approved by the Animal Care and Use Committees overseeing animal welfare at the Valley Biosystems, Charles River Laboratories, and Virscio primate facilities.

### Directed Evolution – Intravitreal Injection, Tissue Harvesting, and Tissue Dissection

For each of the 6 rounds of selection, a single male Cynomolgus macaque (*Macaca fascicularis*) age 4-10 years old and weighing at least 4 kg was dosed via intravitreal injection through the sclera (approximately 3 mm behind the limbus). The animal was anesthetized with 15 mg/kg Ketamine IM and given the topical anesthetic Proparacaine. In addition, IM injection of 5 mg/kg Ketofen was used to minimize post-injection discomfort. The library (in a 100 μL volume) was administered to each eye using a 30-gauge needle attached to a one mL syringe. At the conclusion of injections, topical atropine 1% and triple antibiotic ointment with 1% hydrocortisone was applied to the cornea. Once variants reached the target cells within the retina, tissue was harvested. Euthanasia was performed by trained veterinary staff using 100 mg/kg pentobarbital sodium intravenous injection following the completion of in-life. Eyes were enucleated and stored in 4% paraformaldehyde at 4°C until dissection.

Eyes were cut along the ora serrata with a scalpel, and the anterior segment was removed. Relief cuts were made into the retina around the fovea to enable flat mounting of the retina, and the vitreous was removed. Each retina was processed such that multiple peripheral, medial, and central samples of the ganglion cell layer, inner nuclear layer, outer nuclear layer, and retinal pigment epithelial layer could be obtained from across the retina. Six samples of the retina from each quadrant (superior, inferior, nasal, and temporal) were collected. Samples were rinsed with PBS, then incubated in 30% sucrose. DNA was isolated, and AAV genomic DNA was amplified and recovered by PCR amplification from each of these samples. The process was repeated for a total of 6 rounds of selection. All studies were performed in accordance with regulations outlined in the USDA Animal Welfare Act (9 CFR, parts 1, 2, and 3) and the conditions specified by the National Research Council (2011). Research was conducted under an Institutional Animal Care and Use Committee (IACUC) approved protocol at Valley Biosystems, a fully AAALAC (Association for the Assessment and Accreditation of Laboratory Animal Care) accredited Contract Research Organization.

### Directed Evolution – Viral Genome Amplification and Sequencing Analysis

DNA was isolated from the RPE and retina sections described above using the QIAamp DNA Mini isolation kit (Qiagen). AAV variant *cap* genes were amplified from the tissue by PCR using 5’-GACGTCAGACGCGGAAGCTTC-3’ and 5’-GATTAACAAGCGGCCGCAATTAC-3’ as forward and reverse primers, respectively. The *cap* genes were inserted into the pSub2 library packaging plasmid using NotI and HindIII, as previously described. ^48, 49, 51, 53, 54^

*Cap* genes were then sequenced by third-party DNA sequencing facilities. The sequencing files were analyzed using Geneious software (Biomatters). Peptide homology was assessed using the protein-protein Basic Local Alignment Search Tool (BLAST). A three-dimensional model of the R100 capsid was rendered in Pymol (DeLano Scientific) using the AAV2 crystal structure (Protein Databank accession number 1LP3).

### AAV Manufacturing

Recombinant R100 and AAV2 viral vectors were produced by transient transfection in HEK293 cells. Cells were cultured in DMEM supplemented with FBS and were maintained at 37°C in a 5% CO_2_ environment. Cells were triply transfected (payload, capsid, and helper plasmids) using polyethylenimine (PEI). At times 48 to 96 hours post-transfection, viral particles were harvested from cells and/or supernatant and cells lysed via microfluidization. Cell lysate and/or supernatant was enzymatically treated to degrade plasmid and host-cell DNA, then clarified and concentrated by tangential flow filtration (TFF). The TFF retentate was then loaded onto an affinity resin column for purification. Following pH-gradient elution, post-affinity material was buffer exchanged, then further purified (if needed) by anion-exchange chromatography. Purified rAAV was then formulated into DPBS with 0.001% polysorbate-20, sterile filtered, and filled to yield rAAV Drug Product.

### Neutralizing Antibody Assay

2v6.11 cells were plated at a density of 3×10^4^ cells/well in 96 well plates 24 hours prior to infection. rAAV vectors encoding firefly luciferase driven by the CAG promoter were incubated at 37°C for 1 hour with individual serum samples prior to infection, and cells were then infected at a genomic MOI of 1,000. Luciferase activity was assessed 48 hours post infection using the Luc-Screen Extended-Glow Luciferase Reporter Gene Assay System (Invitrogen) or the ONE-Glo Luciferase Assay System (Promega) and quantified using the BioTek Cytation 3 Cell Imaging Multi-Mode Reader and Gen5 software.

Prior to enrollment in studies, non-human primates (NHP) serum was screened for the presence of neutralizing antibodies against AAV1, AAV2, AAV5, and AAV8 (for directed evolution) or R100 (for characterization). NHPs were enrolled in studies when samples resulted in less than 50% neutralization of AAV transduction at a 1:10 serum dilution.

### Generation of Normal and Patient-Derived Induced Pluripotent Stem Cells

The following cell lines were obtained from the NIGMS Human Genetic Cell Repository at the Coriell Institute for Medical Research: GM25421, GM25383. These cells were approved for reprogramming use through a Coriell Institute Statement of Research Intent Form.

Cellular reprogramming of normal fibroblasts and fibroblasts from Choroideremia patients, referred to as CHM1 and CHM2 (Coriell Institute; patient ID: GM25421, GM25383, respectively), was performed by Simplicon RNA reprogramming (EMD Millipore) using synthetic *in vitro* transcribed RNA expressing four reprogramming factors (OKS-iG; Oct4, Klf4, Sox2 and Glis1) in a polycistronic transcript that self-replicates for a limited number of cell divisions.

At day 10, approximately 5×10^4^ –1×10^5^ reprogrammed cells were re-plated on growth factor reduced Matrigel (Corning) in mouse embryonic fibroblasts (MEF)-conditioned medium containing B18R protein (200 ng/mL) supplemented with human iPSC Reprogramming Boost Supplement II (EMD Millipore). At day 20, reprogrammed cells, recognized by altered morphology and ability to form small colonies, were transitioned to mTeSR-1 media (Stem Cell Technologies). Colonies of approximately 200 cells or larger were isolated manually and plated on growth factor reduced Matrigel coated plates in mTeSR-1 medium. Normal FB-iPSC and CHM-iPSC lines were expanded from a single colony. All iPSC lines were cultured on Vitronectin XF (Stem Cell Technologies) in mTeSR-1 maintenance medium and sub-cultured using Gentle Cell Dissociation Reagent (Stem Cell Technologies), every four to five days at 70-80% confluence. To ensure tri-lineage differentiation capacity, embryoid bodies (EBs) were formed in suspension culture for one week and then differentiated in adherent conditions for an additional four weeks in mTeSR-1 basal medium, plus 20% Knockout Serum Replacement (Thermo Fisher Scientific). Germ layer differentiation was assayed by immunocytochemistry for ectoderm, mesoderm, and endoderm markers.

### Generation of Human Retinal Pigmented Epithelial (RPE) Cells

RPE cells were generated from all iPSC lines by a directed differentiation protocol as previously described.^83, 84^ Briefly, iPSCs were passaged directly onto Matrigel-coated plates (BD Biosciences) in DMEM/F12 with 1× B27, 1× N2, and 1× NEAA (Invitrogen). From days 0 to 2, 50 ng/ml Noggin, 10 ng/ml Dkk1, 10 ng/ml IGF1 (R&D Systems Inc.), and 10 mM nicotinamide (Sigma-Aldrich) were added to the base medium. From days 2 to 4, 10 ng/ml Noggin, 10 ng/ml Dkk1, 10 ng/ml IGF1, 5 ng/ml bFGF, and 10 mM nicotinamide were added to the base medium. From days 4 to 6, 10 ng/ml Dkk1, 10 ng/ml IGF1, and 100 ng/ml Activin A (R&D Systems) were added to the base medium. From days 6 to 14, 100 ng/ml Activin A, 10 μM SU5402 (EMD Millipore), and 1 mM VIP (Sigma-Aldrich) were added to the base medium. At day 14, the cells were mechanically enriched by scraping away cells with non-RPE morphology. Subsequently, the remaining RPE were digested using TrypLE Express (Invitrogen) for ∼5 minutes at 37°C. The cells were passed through a 30-μm cell strainer and seeded onto Matrigel-coated tissue culture plastic, transwell membranes (Corning Enterprises), or CC2-treated chambered slides in XVIVO-10 media (Lonza).

RPE cells from the ESI-017 were generated by dual SMAD inhibition with 100 nM LDN 193189 and 10 μM SB431542 (Stemgent) in combination with 10 mM of another neural inducer, nicotinamide (Sigma-Aldrich), during the first six days. During this time, Wnt signaling was inhibited by a small molecule inhibitor, 5 μM IWP-2 (R&D Biosystems). From day six to 14, cultures were subjected to 10 ng/mL BMP4, 100 ng/mL Activin A (R&D Biosystems), and 3 μM CHIR (R&D System Inc.). Culture media used for the first 14 days was composed on Advanced DMEM/F12 supplemented with 1× N2 and 1× B27 (ThermoFisher). The RPE cells were manually enriched by removing cells with non-RPE morphology. Subsequently, the remaining immature RPE cells were digested using TrypLE Express (ThermoFisher) for 5 minutes at 37°C. The cells were passed through a 30 μm cell strainer and seeded onto Matrigel matrix (Corning) coated plates and maintained in XVIVO-10 media (Lonza). The medium was changed every two to three days. Every 30 days, the cells were harvested using TrypLE Express and re-plated at a density of 100,000 cells per cm^2^ in the presence of 10 µM Y-27632 (Stem Cell Technologies) for the first 14 days. For experiments, RPE cells were passaged and seeded on Vitronectin XF (Stem Cell Technologies)-coated plates according to the manufacturer’s instructions. Following 45 days of development, over 90% of cells expressed mature RPE-specific markers, including RPE65, BEST1, and PMEL17, synthesized VEGF and PEDF, and were capable of phagocytosing rod outer segments (Supplemental Figure S2).

### Functional Characterization of Human Retinal Pigment Epithelial Cells

To perform the rod outer segment phagocytosis assay, RPE cells were cultured using a formulated medium.^85^ All cells were plated in quadruplicate at 100,000 cells per cm^2^ in 0.1% gelatin-coated black-walled, clear bottom 96 well plates and cultured for 30 days. Photoreceptor rod outer segments (ROS) were isolated from bovine eyes (Sierra for Medical Science, Whittier, CA) as previously described^86^ and fluorescently labeled with fluorescein isothiocyanate (FITC) protein (ThermoFisher). In some conditions, cultured cells were treated with 62.5 μg/mL αVβ5 integrin function-blocking antibody (Abcam, Cambridge, UK) or IgG isotype control (Abcam) for 30 minutes at 37°C. Following the initial antibody incubation, cells were challenged with 1×10^6^ FITC-ROS per well for five hours at 37°C and 5% CO_2_.^83, 87^ After ROS incubation, the wells were washed six times with PBS, and 0.4% trypan blue was added for 20 minutes to quench fluorescence from extracellular ROS. Each well was imaged using epifluorescent microscopy, and integrated pixel density of photomicrographs was determined with Image J software using a rolling pixel radius of 50 (National Institutes of Health, Bethesda, MD).

To measure levels of Pigment Epithelium-Derived Factor (PEDF) and Vascular Endothelial Growth Factor (VEGF) in RPE cells, conditioned media was collected from RPE cells after 48 hours of culture, and ELISA was performed (PEDF, Express Bio/MD Bioproducts; VEGF, R&D Systems Inc.) in accordance with the manufacturer’s instructions.

### Generation of Human Retinal Ganglion Cells

RGCs were generated using established protocols following in-house optimization.^57, 58^ Embryoid bodies were created from Fibroblasts-iPSCs and transitioned into Neural Induction Medium (NIM), containing N2 supplement. After seven days, EBs were seeded at a high density into 6-well tissue culture plates. NIM was changed every other day until a retinal phenotype was observed. Between days 28-42, retinal centers were dissected and cultured as spheres in Retinal Differentiation Medium (RDM) containing B27 supplement without retinoic acid. Optic vesicles displaying a retinal phenotype were manually isolated and seeded onto poly-L-ornithine and laminin coated plates between days 40-42. At day 42, RDM was supplemented with 10% fetal bovine serum (GE LifeSciences, Pittsburgh, PA) and 100 µM taurine (Sigma Aldrich, St-Louis, MO). Cells were cultured until days 60-70 when RGC marker expression (Brn3a) appeared, media was changed every two to three days. Following 60-70 days of differentiation, expression of Brn3a, a marker for RGCs, was detected primarily in adhered neural spheres that retained 3D structure. Further differentiation led to the emergence of photoreceptors and decline of the RGC population, as neural spheres expanded outward to from a monolayer.

### Generation of Human Photoreceptors

To develop self-organized retinal aggregates referred to as “optic cups”, hPSCs were dissociated to form small clusters and cultured in suspension to generate embryoid bodies (EBs).^88^ EBs were slowly transitioned from mTeSR-1 medium to neural induction medium (NIM) containing advanced DMEM/F12, 1× N2 (ThermoFisher) and 2 µg/mL Heparin (Sigma-Aldrich) at the following mTeSR-1:NIM ratios, Day 0 – 3:1 (supplemented with 10 µM Y-27632, Stem Cell Technologies), Day 1 – 1:1, Day 2 – 1:3, Day 3 – complete NIM. After seven days of differentiation, EBs were plated in the presence of 10% fetal bovine serum (ThermoFisher), for the first 24 hours to enhance cell adhesion. Medium was changed every other day until day 24 of differentiation. At day 24, the center of colonies containing retinal progenitors were mechanically dislodged and maintained in Retinal Differentiation Medium (RDM) composed of 1:1 ratio of DMEM/F12 and Advanced DMEM medium with 1× B27 supplement (ThermoFisher), and 1% penicillin/streptomycin (ThermoFisher) in non-tissue culture treated 60 mm dishes to allow the development of a three-dimensional (3D) optic cup-like structures. The media in neural progenitor cultures was changed every two to three days. By day 30-35 of differentiation, retinal neurospheres, as recognized by a bright appearance with a surrounding golden ring, were manually selected and isolated from non-retinal dark neurospheres. Media was replaced every two to three days until day 48 of differentiation. At day 48, retinal spheres were plated onto Poly-L-ornithine (Sigma-Aldrich)/laminin (ThermoFisher) coated plates to generate two-dimensional, adherent mature photoreceptors. Cultures were maintained in RDM supplemented with 10 µM DAPT (R&D Systems Inc.), exchanging half of the media every other day until the desired stage of differentiation was reached.

For generation of optic cups in 3D suspension culture, retinal progenitors were maintained in suspension for over 120 days in RDM in the presence of 10 µM DAPT. Half of the media was changed every two days. Following differentiation, cultures expressed photoreceptor markers including recoverin, rhodopsin, S-opsin, and M/L-opsin (Supplemental Figure S2).

### Transduction of Human Retinal Cells by AAV Vectors

After human RPE cells reached maturity at 30 days in culture, AAV vectors R100.CAG-EGFP, AAV2.CAG-EGFP, 4D-110, or 4D-140 were administered to the cell culture medium. For determination of VEGF and anti-VEGF levels, the media was changed after 72 hours and collected for assay 96 hours later. For transduction of RGCs, R100.CAG-EGFP or 4D-140 were administered to the cell culture medium at various MOI (viral genomes per cell): 10,000 (R100.CAG-EGFP) and 0.1, 1, 10, 100, and 1,000 (4D-140). Media was changed two days following transduction and subsequently changed every three days. Media was collected for analysis 12-15 days following transduction to assay for secreted VEGF and anti-VEGF scFv. For transduction of photoreceptors, R100.CAG-EGFP was applied to adherent culture medium (for 2D cultures) or cell suspension medium (for optic cups in 3D cultures). Viral particles were removed from media and replaced with fresh media after 48 hours.

### VEGF and Anti-VEGF ELISA

Media from transduced RPE and RGCs was assayed for free VEGF protein (not bound by anti-VEGF) with the Quantikine VEGF ELISA kit (R&D Systems Cat# DVE00). Media and aqueous fluid from NHPs were assayed for free anti-VEGF scFv (not bound to VEGF) with the Aflibercept ELISA kit (Eagle Biosciences, Immunoguide IG-AA115) using HRP-conjugated Protein L (Pierce/ThermoFisher Cat# 32420) as the detection reagent. The concentration of free anti-VEGF scFv was calculated relative to a purified protein standard initially expressed from HEK293E cells (GenScript).

### Immunocytochemistry and Immunofluorescence

Cells were fixed with 4% paraformaldehyde (PFA; Santa Cruz Biotechnology, Dallas, TX) for 15 minutes at 4°C. Cells were incubated with primary and secondary antibodies (Supplemental Table 1) in a solution of PBS with 0.2% Triton-X100 (Sigma-Aldrich), 2% bovine serum albumin (Calbiochem), and 5% goat serum. Primary antibodies were incubated overnight at 4°C. Secondary antibodies were incubated for one hour at room temperature. Cells were counterstained with DAPI (Sigma Aldrich) in PBS for five minutes at room temperature.

Optic cups were fixed in 4% paraformaldehyde at 4°C overnight, then subjected to a stepwise incubation with 15% and 30% sucrose at 4°C overnight. Optic cups were embedded in molds using optimum cutting temperature solution (VWR) and cut in 20 μm sections using a −20°C cryostat and stored at −80°C.

Cells were imaged using a Zeiss Axio Observer.D1 fluorescent microscope (Carl Zeiss Microscopy LLC, White Plains, NY). Image processing was performed using Zeiss Zen 2 software and FIJI.

### Flow cytometry

Cells were fixed and stained using the Fixation/Permeabilization kit (BD Biosciences) according to manufacturer’s protocol. Flow cytometry was performed using the Accuri system (BD Biosciences).

### R100 Vector and 4D-140 Characterization – Intravitreal Injection

For the R100 vector characterization studies, male and female cynomolgus macaques (*Macaca fascicularis*) aged 2-8 were dosed via two 50 µL intravitreal injections into each eye through the sclera (approximately 3 mm behind the limbus) for a total dose volume of 100 µL/eye. Doses of 1×10^11^ vg/eye and 1×10^12^ vg/eye were evaluated. The animals were anesthetized with 10 mg/kg Ketamine IM and given the topical Proparacaine, Tropicamide, and Phenylepherine ophthalmic solutions to eliminate pain or discomfort. In addition, IM injection of 40-80 mg methylprednisolone was administered weekly post-injection. Euthanasia was performed by trained veterinary staff using by intravenous injection of euthanasia solution at defined study termination times. Eyes were enucleated and fixed as described below. All studies were performed in accordance with regulations outlined in the USDA Animal Welfare Act (9 CFR, parts 1, 2, and 3) and the conditions specified by the National Research Council (2011). Research was conducted under an Institutional Animal Care and Use Committee (IACUC) approved protocol at Charles River Laboratories, a fully AAALAC (Association for the Assessment and Accreditation of Laboratory Animal Care) accredited Contract Research Organization.

For the 4D-140 study, male and female African green monkeys (*Chlorocebus sabaeus*) aged 4-13 years old received bilateral (OU) IVT administrations each consisting of two 50 µL volumes of 4D-140 or vehicle. IVT doses were administered under local anesthesia (0.5% proparacaine) using a 31G 5/16-inch needle (Ulticare Vet RX U-100, # 09436) ∼2.5 mm posterior to the limbus. Animals were anesthetized for all procedures and ophthalmic evaluations (8.0 mg/kg ketamine/1.6 mg/kg xylazine, intramuscularly [IM] to effect). General well-being was assessed before, during, and after sedation as well as twice daily on non-procedure days. This study was performed in accordance with the ARVO Statement for the Use of Animals in Ophthalmic and Vision Research. Research was conducted under an Institutional Animal Care and Use Committee (IACUC) approved protocol at Virisco, a fully AAALAC (Association for the Assessment and Accreditation of Laboratory Animal Care) accredited Contract Research Organization.

### Vector Characterization – Clinical Examination and In-Life Imaging

Ophthalmoscopic examinations and ocular imaging were conducted by a board-certified veterinary ophthalmologist pre-administration and at various timepoints post-administration. Examinations included indirect ophthalmoscopy and slit-lamp biomicroscopy. Wide-field, color fundus imaging (RetCam Shuttle, Clarity Medical Systems) was conducted on all animals pretest and at various timepoints post-administration. Confocal scanning laser ophthalmoscopy (cSLO) imaging (Spectralis, Heidelberg Engineering) was conducted on all animals at various timepoints post-administration and consisted of infrared reflectance (IR) and blue autofluorescence (BAF). Optical coherence tomography (OCT) imaging (Spectralis, Heidelberg Engineering) was conducted at various timepoints post-administration. OCT imaging consisted of a single, horizontal, high-resolution line scan from optic nerve head through fovea.

### Vector Characterization –Tissue Harvesting

An approximately 6 mm biopsy punch was made in the middle of the cornea and lens of each enucleated cynomolgus macaque eye. The eyes were then fixed in 4% paraformaldehyde for 4 hours at 4°C and stored in 30% sucrose in PBS^-/-^ with 0.05% sodium azide at 4°C. Lens and cornea tissue were removed by creating an incision approximately 10 mm diameter around the lens closest to ciliary body. Once the lens and cornea were removed, 4 relief cuts of approximately 1 cm from anterior to posterior were made to allow the retinal tissue to lay flat. The retina was then further dissected into petals encompassing the superior, inferior, temporal, nasal, and optic nerve/fovea/macula (ON/FOV) regions. The petals were mounted in OCT in individual plastic molds. Samples were then frozen on dry ice, partially submerged in an ethanol bath. Mounted tissues were then stored at −20°C until sectioning.

### Histology

Prior to sectioning, mounted eyes were equilibrated in the cryostat chamber at −20°C for a minimum of 30 minutes. Each petal was then mounted individually to the chuck using additional OCT with the left side facing upwards. Sagittal sections 20 µm thick were then generated through the eye. Each section was placed onto warmed positively-charged slides. Each eye produced around 20 slides with seven sections per slide. Each slide represented either nasal, temporal, superior, inferior, or ON/FOV regions of the retina. Slides were allowed to dry at room temperature for one hour and then stored in slide boxes at −20°C.

Retinal tissue is known to possess a high level of autofluorescence detected at the 488 nm wavelength. Because of these challenges, an anti-GFP antibody was used in conjunction with direct GFP expression to visualize GFP in all cynomolgus macaque retinal sections.

Slides were removed from −20°C and allowed to dry for 20 minutes at room temperature. Rubber cement was used to outline sections and to create a barrier to allow fluid to stay on the slide. Rubber cement was allowed to dry fully for 20 minutes. Slides were then washed in 300 µl PBS per slide for 5 minutes each, 3 times. The slides were permeabilized and blocked in blocking buffer comprised of PBS with 0.5% Triton X-100, and 10% goat serum for one hour.

Slides were then incubated with primary antibodies (Supplemental Table 1) in blocking buffer at 4°C overnight in staining chambers. Slides were then washed with three 5 minute PBS + 0.5% Triton X-100 washes. Slides were then incubated with Alexa Fluor-conjugated secondary antibodies (Supplemental Table 1) for one hour at room temperature in blocking buffer. Slides were then washed three times with PBS for 5 minutes per wash. Tissues were counterstained with DAPI nuclear stain for 20 minutes. DAPI was washed out with PBS three times 5 minutes each. Slides were mounted using Prolong gold and cover slips were sealed with clear nail polish.

### Imaging and Image Processing

Images were acquired using a Zeiss Axio Observer Z1 microscope or a Zeiss Axio Observer D1 microscope using a 10×, 20×, and 63× objectives. Images were taken with brightfield, DAPI, GFP, dsRED, and Cy5 filters to collect phase contrast, DAPI, GFP, Alexa Fluor 555, and Alexa Fluor 680, respectively. For tiling, sequential multi-channel images with approximately 5% overlap were acquired.

Images were processed using Zen Blue 2.3 software and FIJI Image J software. Channels were adjusted for brightness and contrast. Color merges were made with all fluorescence channels and added to the end of the stack to generate a stack montage. For tiling of images, the “grid/collection stitching” plugin was used. Images were cropped and rotated to have the superior part of the eye on top.

### Histological Scoring of GFP Expression

In order to quantify the extent of variant transduction and subsequent protein expression in multiple cell layers across large portions of the cynomolgus macaque retina, a histological scoring method was developed. Scoring of GFP expression was performed using a Zeiss Axio Observer D1 microscope with a 20× objective. All scoring was performed by evaluating fluorescence in the far red channel resulting from an anti-GFP antibody and a fluorescent conjugated secondary antibody. The same exposure setting was used for all scoring. The background fluorescent pixel intensity in the 647 nm channel for each cell layer within each tissue section was determined. Cells within each cell layer are considered positive if the fluorescent pixel intensity within the cell is at least 1.5-fold higher than the background pixel intensity. The RPE and PR layers are scored based on the number of fluorescent cells within the layer. The RGC layer is scored based on the volume of fluorescence within the layer in order to assess both the RGC cell bodies and the accompanying fiber bundles.

The amount of fluorescent cells within each layer is quantified as follows:

| Score | Estimated Percent of GFP+ Cells |
| --- | --- |
| 4 | 76% - 100% |
| 3 | 51% - 75% |
| 2 | 26% - 50% |
| 1 | > 0% - 25% |
| 0 | 0% |

### Molecular Modeling

The solution phase peptide modeling package PEP-FOLD3^89^ was used to generate an ensemble of structural predictions for the R100 peptide sequence and its flanking amino acids. The lowest energy conformer that preserved the global architecture of this spike domain near the capsid surface was grafted onto the AAV2 capsid structure using PyMol (version 2)^90^.

### AAV Binding Assay

RPE cells were incubated on ice for 10 minutes prior to application of viral particles for 30 minutes on ice. Unbound virus was removed by washing three times with 500 μL of ice-cold X-VIVO10. DNA was isolated with the DNeasy Blood & Tissue Kit (Qiagen) according to the manufacturer’s instructions. Quantitative PCR was performed using the TaqMan Universal PCR Master Mix (ThermoFisher) with primers designed to the SV40 polyadenylation region (5’-AGCAATAGCATCACAAATTTCACAA-3’ and 5’-CCAGACATGATAAGATACATTGATGAGTT-3’) and compared to DNA standards to determine the amount of viral DNA.

### Prenylation Assay

The prenylation assay was performed using RPE cell lysates from control RPE cells derived from a normal fibroblast cell-derived iPSC line, non-transduced CHM1 and CHM2 RPE cells, or CHM1 and CHM2 RPE cells transduced with 4D-110. Following a wash with PBS^-/-^, cell lysates were prepared in cold Prenylation Buffer (500 µM HEPES pH 7.0; 50 µM NaCl; 2 µM MgCl2; 0.1 µM GDP; 0.5% NP-40; Complete Protease Inhibitor Tablet) and incubated on ice for 10-15 minutes. Protein concentrations were determined using a BCA protein assay (ThermoFisher). Protein concentrations were normalized and lysates adjusted to 2 µM DTT.^91^ Prenylation reactions were performed using 20 µL of lysate corresponding to 50-200 µg protein. The reaction for the functional complex was composed of 2 µM RabGGTase, 4 µM Rab27a and 4 µM BiotinGeranyl-PPi (Jena Bioscience, Jena, Germany). Reactions were incubated at 25°C for 5 hours and stopped by adding 4× Sample Buffer (Bio-Rad, Hercules, CA), DTT to 40 mM and heating at 70°C for 10 minutes. Western blotting was carried out on XT Criterion gels according to manufacturer’s protocols. Prenylation reactions were analyzed using streptavidin-HRP (Abcam).

### Rab27A Trafficking Assay

RPE cells were seeded onto Vitronectin XF coated eight chambered slides at 25,000 cells per cm^2^ in XVIVO-10 media. Two days after seeding, the CHM RPE cells were transduced with 4D-110 at an MOI of 5,000. Fourteen days post infection, cells were fixed analyzed.

### Laser-Induced Choroidal Neovascularization

At 42 days post-dose, six laser spots were placed within perimacular region in each eye by an ophthalmologist employing an Iridex Oculight TX 532 nm laser with a laser duration of 100 ms, spot size 50 μm, power 750 mW. Color fundus photography was performed immediately after the laser treatment to document the laser lesions. Any spots demonstrating severe retinal/subretinal hemorrhage immediately post-laser and not resolving by the time of follow-up examinations were excluded from analyses.

### Color fundus photography (CFP) and fluorescein angiography (FA)

Bilateral color fundus images were captured using a Topcon TRCS0EX retinal camera with Canon 6D digital imaging hardware and New Vision Fundus Image Analysis System software under pupil dilation (10% phenylephrine hydrochloride and 1% cyclopentolate hydrochloride). FA was performed with 10% sodium fluorescein (0.1 mL/kg, IV). FA in oculus dexter (OD) preceded angiography in oculus sinister (OS) by 4–6 hours to allow washout of the fluorescein between the angiogram image series (0, 2, 15, 20, 25, 30, 40, and 50 s, 1, 2, 3 and 6 min). Graded scoring of angiograms was performed on fluorescein angiogram series collected at two and four weeks post-laser by a masked investigator. Fluorescein leakage was graded by a masked investigator using the following grading scale: I, no hyperfluorescence; II, hyperfluorescence without leakage and no significant residual staining in late-phase angiograms; III, hyperfluorescence early or mid-transit with late leakage and significant residual staining; IV, hyperfluorescence early or mid-transit with late leakage extending beyond the borders of the treated area.

## Acknowledgements

The authors thank Katie Sullivan, Devi Khoday, and Thomas Mason for generation and analytical release assays of virus lots used in the experiments; Kathy Delaria for technical assistance performing heparin column binding; Jon Melnick for technical assistance generating molecular models of peptide insertion; Elizabeth Alcamo for help with immunofluorescence method development; Gabriel Bolender for coordination of the NHP CNV study; and F. Hoffmann-La Roche Ltd. for financial and scientific support.

## Author Contributions

<u>Melissa Kotterman</u>: conceived and planned directed evolution screen (Figure 1); designed and generated libraries; conceived and planned experiments (Figure 1, Figure 3, Figure 5); analyzed data (Figure 1, Figure 3, Figure 5, Figure S1, Figure S5); generated images (all Figures) wrote the manuscript

<u>Ghezal Beliakoff</u>: performed directed evolution screening (Figure 1); performed retina sectioning, staining, and scoring (Figure 3c, Figure S5); performed retina histology imaging (Figure 3b, d-h); performed cell transduction mechanism experiments; performed retina histology mechanism staining and imaging

<u>Roxanne Croze</u>: derivation of RPE lines, generation of RPE banks, and RPE cell and vector characterization (Figure 2 & Figure S2); performed 2D and 3D culture photoreceptor cell differentiations and vector characterization (Figure 2e & Figure S2); performed derivation of RGCs, generation of banks, and vector characterization (Figure 2d); performed *in vitro* anti-VEGF expression and activity assays in RGC cells (Figure 6f)

<u>Tandis Vazin</u>: conceived and planned experiments (Figure 2, Figure 6); designed human retinal cell differentiation paradigms and characterization assays (Figure 2, Figure 6, Figure S2, Figure S6); wrote portions of the manuscript

<u>Christopher Schmitt</u>: characterization and differentiation of RPE; developed methods for flow and ICC of RPE (Figure 2); analyzed data (Figure 3c, Figure S5); optimized IHC and performed retina sectioning, staining, and scoring (Figure 3c, Figure S5); performed retina histology imaging (Figure 5g-h); characterized transduction efficiency (Figure 2a-c); imaging for RPE and photoreceptors (Figure 2)

<u>Paul Szymanski</u>: designed and generated anti-VEGF therapeutic payloads; optimized anti-VEGF expression and activity assays; performed *in vitro* anti-VEGF expression and activity assays in RPE cells (Figure 6e); performed *in vivo* anti-VEGF expression assays in aqueous fluid (Figure 6g); wrote portions of the manuscript

<u>Meredith Leong</u>: developed protocols and analyzed data for NHP in-life study (Figure 6g); wrote portions of the manuscript

<u>Melissa Quezada</u>: performed and developed, and optimized process for 1 round of directed evolution screening; performed neutralizing antibody assay for NHP enrollment in studies

<u>Jenny Holt</u>: project manager for retina program; coordinated CRO activities for in-life NHP studies

<u>Katy Barglow</u>: generation of molecular models of peptide insertion (Figure 5a-b); planned heparin column binding experiment (Figure 5c); pilot plant supervisor for 2 virus lots used for NHP studies

<u>Mohammad Hassanipour</u>: generated RPE banks (Figure 2, Figure 6, Figure S2, Figure S6); performed capsid binding to RPE experiment (Figure 5f); characterized vector transduction efficiency (Figure 2a-c); 2D culture photoreceptor cells and vector characterization (Figure 2d)

<u>David Schaffer</u>: conceived directed evolution screen; consulted on directed evolution screen design

<u>Peter Francis</u>: developed protocols for NHP in-life studies (Figure 4, Figure S3, Figure S4); consulted on data package for IHC (Figure 3); reviewed/edited manuscript

<u>David Kirn</u>: reviewed/edited manuscript

All co-authors read the manuscript.

## Data & Material Availability Statement

All requests for raw and analyzed data and materials will be promptly reviewed by 4D Molecular Therapeutics to determine if the request is subject to any intellectual property or confidentiality obligations. Any data and materials that can be shared will be released via a material transfer agreement.

## Human Stem Cell Ethics Statement

Embryonic stem cell line 017 was purchased from ESI-BIO (AgeX Therapeutics) and were cultured according to their guidelines. All ESI-BIO cell lines are backed with donor history and testing information in best compliance with current Good Tissue Practice (cGTP) and conform to Global Ethical Standards and Clinical Cell Regulations. Please refer to ESI-BIO’s polices for additional information regarding licensing, consent, and conditions for donations. All induced pluripotent stem cell lines were purchased through Coriell Institute and were cultured according to their instructions. Coriell laboratories operates under ISO 9001:2015 certified standard operating procedures. Informed consent was obtained according to their procedures and detailed in the certificate of analyses. Please refer to Coriell Institutes’ polices for additional information regarding licensing, consent, and conditions for donations.

## Supplemental Data

**Supplemental Figure S1. Frequency of variants within directed evolution sequencing analysis. a**, Round 3 sequencing analysis. **b**, Round 4 sequencing analysis. **c**, Round 5 sequencing analysis. **d**, Round 6 sequencing analysis.

**Supplemental Figure S2: Differentiation and characterization of human pluripotent stem cell derived-retinal cells.** Images illustrating restriction of **a**, hESC and **b**, iPSC lines to an RPE phenotype after 45 days of differentiation by immunocytochemical analysis using antibodies against transcriptional factors Melanogenesis Associated Transcription Factor (MITF) and Orthodenticle Homeobox 2 (OTX2), as well as other RPE cell markers including zonula occludens (ZO-1) and Bestrophin 1 (BEST1). The nuclei were counterstained with DAPI (blue). **c**, levels of photoreceptor outer segment (ROS) internalization via phagocytosis measured by pixel density of fluorescence micrographs in the presence or absence of αVβ5 integrin function-blocking antibody or IgG isotype control. All quantitative measurements were carried out in n = 3; data presented as mean ± s.d.; * p < 0.05, compared to the “No ROS” condition; † p < 0.05, compared to the ROS condition; two-tailed t-test. Bar graphs illustrating levels of secreted **d**, vascular endothelial growth factor (VEGF) and **e**, pigment epithelium-derived factor (PEDF) from hESC- and iPSC-derived 30-day RPE cells during a 48 hour period as compared to human hepatocytic carcinoma HepG2 and human embryonic kidney 293T (HEK293T) control cell lines. **f**, Schematic representation of the differentiation paradigm used to generate photoreceptors in adherent cultures or in suspension in optic vesicles from human pluripotent stem cells. **g**, Representative brightfield image of iPSC-derived colonies containing retinal centers after 24 days of differentiation. **h**, Representative images showing identification of (left) optic vesicle-derived rod (rhodopsin; red) and cone (M/L Opsin; green) photoreceptors and (right) optic vesicle-derived rod (rhodopsin; red) and cone (S Opsin; green) photoreceptors after 192 days of differentiation.

**Supplemental Figure S3. Representative fundus fluorescence images for NHP dosed in three separate studies. a**, NHPs received a dose of 1×10^12^ vg/eye of R100.CAG-EGFP. Top row (left to right): 2 weeks post-administration, NHP ID 304 right eye, NHP ID 304 left eye, NHP ID 305 right eye. Second row (left to right): 3 weeks post-administration, NHP ID 201 right eye, NHP ID 201 left eye, NHP ID 205 right eye, NHP ID 205 left eye. Third row (left to right): 16 weeks post-administration, NHP ID 1501 left eye, NHP ID 1502 left eye, NHP ID 1503 left eye. Fourth row (left to right): 16 weeks post-administration, NHP ID 1501 right eye, NHP ID 1502 right eye, NHP ID 1503 right eye. Bottom row (left to right): 26 weeks post-administration, NHP ID 202 right eye, NHP ID 202 left eye, NHP ID 207 right eye, NHP ID 207 left eye. **b**, NHPs received either a dose of 3×10^11^ vg/eye or 1×10^12^ vg/eye of R100.CAG-EGFP. Top panel: 2 weeks post-administration, NHP ID 304 right eye, NHP ID 304 left eye, NHP ID 305 right eye Bottom panel: 3 weeks post-administration, NHP ID 302 left eye, NHP ID 303 left eye, NHP ID 303 right eye

**Supplemental Figure S4. Representative fundus fluorescence images.** NHP ID 1501 left eye received a dose of 1×10^12^ vg/eye of R100.CAG-EGFP. Top row (left to right): week 1, week 3, week 5; middle row (left to right): week 7, week 8, week 9; bottom row (left to right): week 11, week 13, week 16.

**Supplemental Figure S5. IHC Scoring Heat Map Visualization.** Each eye received a dose of 1×10^12^ vg/eye of R100.CAG-EGFP. **a**, NHP ID 304 right eye. **b**, NHP ID 304 left eye. **c**, NHP ID 305 right eye. **d**, NHP ID 302 left eye. **e**, NHP ID 303 left eye. **f**, NHP ID 303 right eye.

**Supplemental Figure S6. Characterization of choroideremia iPSC lines and retinal pigment epithelial cells.** Immunocytochemical analysis using antibodies against pluripotency transcriptional factors NANOG (orange), OCT4 (green), and SOX2 (red), in **a**, CHM1, and **b**, CHM2 iPSC lines. Representative images of **c**, CHM1, and **d**, CHM2 cultures randomly differentiated into ectodermal, mesodermal, and endodermal cell lineages as indicated by the expression of TUJ1 (red), α-smooth muscle actin (ASMA; red), and Hepatocyte Nuclear Factor 4 Alpha (HNF4A; green), respectively. Images illustrating restriction of **e**, the CHM1, and **f**, CHM2 iPSC lines, to an RPE phenotype after 45 days of differentiation by immunocytochemical analysis using antibodies against transcriptional factors Melanogenesis Associated Transcription Factor (MITF) and Orthodenticle Homeobox 2 (OTX2), as well as other RPE cell markers including RPE65 and Zonula Occludens (ZO-1). The nuclei were counterstained with DAPI. Scale bar = 100 µm. **g**, Levels of photoreceptor outer segment (ROS) internalization via phagocytosis measured by pixel density of fluorescence micrographs in the presence or absence of αVβ5 integrin function-blocking antibody or IgG isotype control. All quantitative measurements were carried out in n = 3; data presented as mean ± s.d.; * p < 0.05, compared to the “No ROS” condition; † p < 0.05, compared to the ROS condition; two-tailed t-test. **h**, Gel images illustrating the level of REP1 protein by western blot analysis and incorporation of a biotinylated prenyl donor as a measure of prenylation in cell lysates from untransduced cells and 4D-110 transduced cells. CHM2 RPE cells as compared to normal iPSC-derived RPE cells. Bar graphs illustrate the average band intensity of biotinylated RAB27A relative to the housekeeping protein GAPDH of each sample. All quantitative measurements were carried out in n = 3; data presented as mean ± s.d.; * p < 0.05, compared to untransduced CHM RPE; two-tailed t-test.

**Supplemental Figure S7. Safety of 4D-140 Following Intravitreal Administration. a**, quantification of intraocular pressure (IOP); data presented as mean ± s.d.. **b**, aqueous cells **c**, aqueous flare and **d**, vitreous cells observed in individual NHP eyes by slit lamp microscopy. **e**, Summary of ocular inflammatory responses observed, grouped based on dose of 4D-140 administered. Scores are shown as mean ± s.e.m.; n = 12 eyes (n = 6 NHP) per group. AC = aqueous cells; AF = aqueous flare; KP = keratic precipitates; LCD = lens capsule deposits VC = vitreous cells.

**Supplemental Figure S8. Example Gating Strategy for Flow Cytometry. a**, ancestry gates for representative sample from flow experiments in Figures 2A & 2B. **b**, ancestry gates for representative sample from flow experiments in Figure 2C. **c**, ancestry gates for representative sample from flow experiments in Figures 5D & 5E.

**Supplemental Figure S9. R100 Capsid Protein Sequence.** Capsid protein sequence alignment comparing R100 and parental AAV2.

**Supplemental Table 1.**
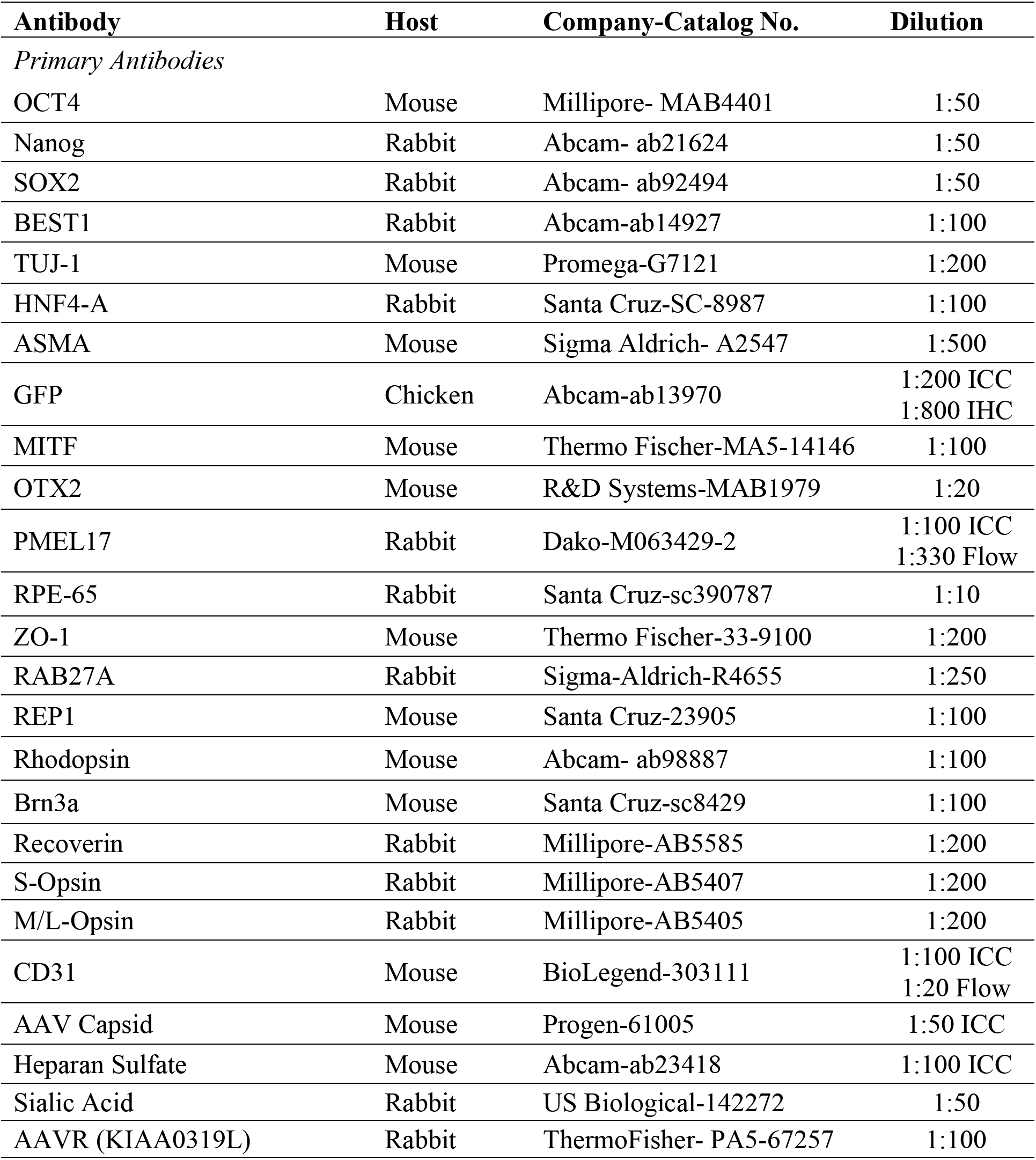
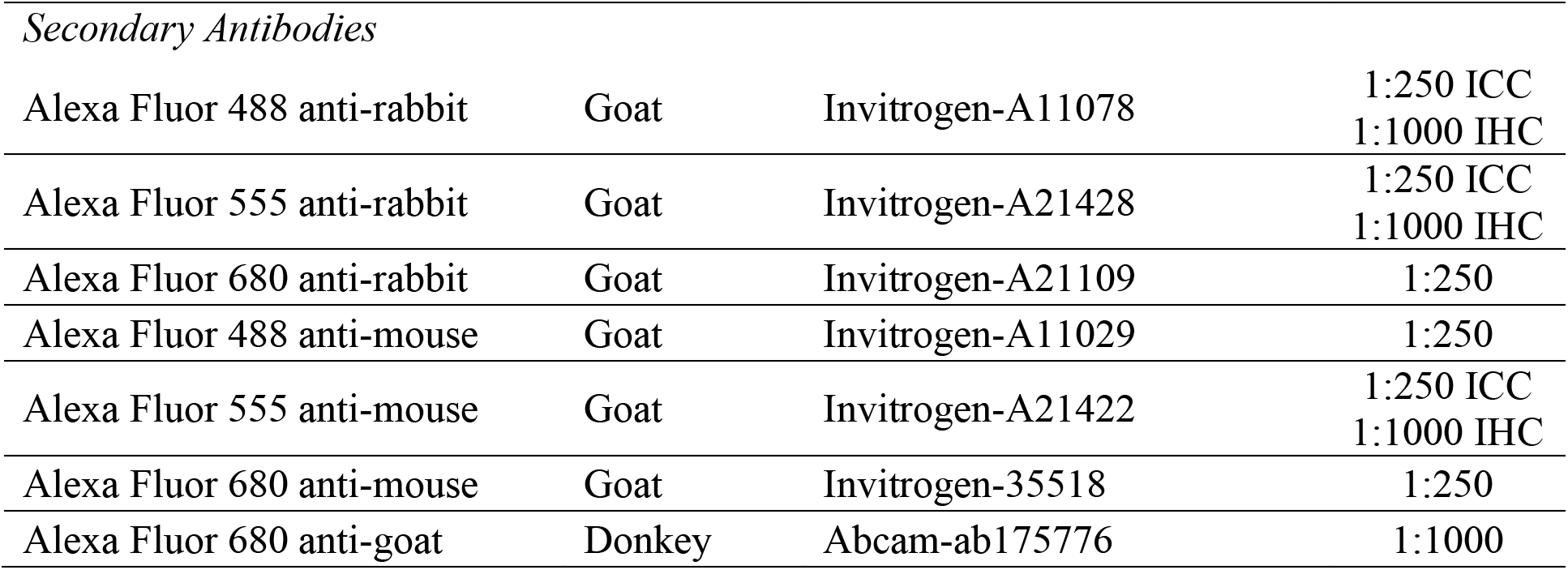

| **Antibody** | **Host** | **Company-Catalog No.** | **Dilution** |
| --- | --- | --- | --- |
| *Primary Antibodies* |  |  |  |
| OCT4 | Mouse | Millipore- MAB4401 | 1:50 |
| Nanog | Rabbit | Abcam- ab21624 | 1:50 |
| SOX2 | Rabbit | Abcam- ab92494 | 1:50 |
| BEST1 | Rabbit | Abcam-ab14927 | 1:100 |
| TUJ-1 | Mouse | Promega-G7121 | 1:200 |
| HNF4-A | Rabbit | Santa Cruz-SC-8987 | 1:100 |
| ASMA | Mouse | Sigma Aldrich- A2547 | 1:500 |
| GFP | Chicken | Abcam-ab13970 | 1:200 ICC<br>1:800 IHC |
| MITF | Mouse | Thermo Fischer-MA5-14146 | 1:100 |
| OTX2 | Mouse | R&D Systems-MAB1979 | 1:20 |
| PMEL17 | Rabbit | Dako-M063429-2 | 1:100 ICC<br>1:330 Flow |
| RPE-65 | Rabbit | Santa Cruz-sc390787 | 1:10 |
| ZO-1 | Mouse | Thermo Fischer-33-9100 | 1:200 |
| RAB27A | Rabbit | Sigma-Aldrich-R4655 | 1:250 |
| REP1 | Mouse | Santa Cruz-23905 | 1:100 |
| Rhodopsin | Mouse | Abcam- ab98887 | 1:100 |
| Brn3a | Mouse | Santa Cruz-sc8429 | 1:100 |
| Recoverin | Rabbit | Millipore-AB5585 | 1:200 |
| S-Opsin | Rabbit | Millipore-AB5407 | 1:200 |
| M/L-Opsin | Rabbit | Millipore-AB5405 | 1:200 |
| CD31 | Mouse | BioLegend-303111 | 1:100 ICC<br>1:20 Flow |
| AAV Capsid | Mouse | Progen-61005 | 1:50 ICC |
| Heparan Sulfate | Mouse | Abcam-ab23418 | 1:100 ICC |
| Sialic Acid | Rabbit | US Biological-142272 | 1:50 |
| AAVR (KIAA0319L) | Rabbit | ThermoFisher- PA5-67257 | 1:100 |
| *Secondary Antibodies* |  |  |  |
| Alexa Fluor 488 anti-rabbit | Goat | Invitrogen-A11078 | 1:250 ICC<br>1:1000 IHC |
| Alexa Fluor 555 anti-rabbit | Goat | Invitrogen-A21428 | 1:250 ICC<br>1:1000 IHC |
| Alexa Fluor 680 anti-rabbit | Goat | Invitrogen-A21109 | 1:250 |
| Alexa Fluor 488 anti-mouse | Goat | Invitrogen-A11029 | 1:250 |
| Alexa Fluor 555 anti-mouse | Goat | Invitrogen-A21422 | 1:250 ICC<br>1:1000 IHC |
| Alexa Fluor 680 anti-mouse | Goat | Invitrogen-35518 | 1:250 |
| Alexa Fluor 680 anti-goat | Donkey | Abcam-ab175776 | 1:1000 |

**Supplemental Table 2.**
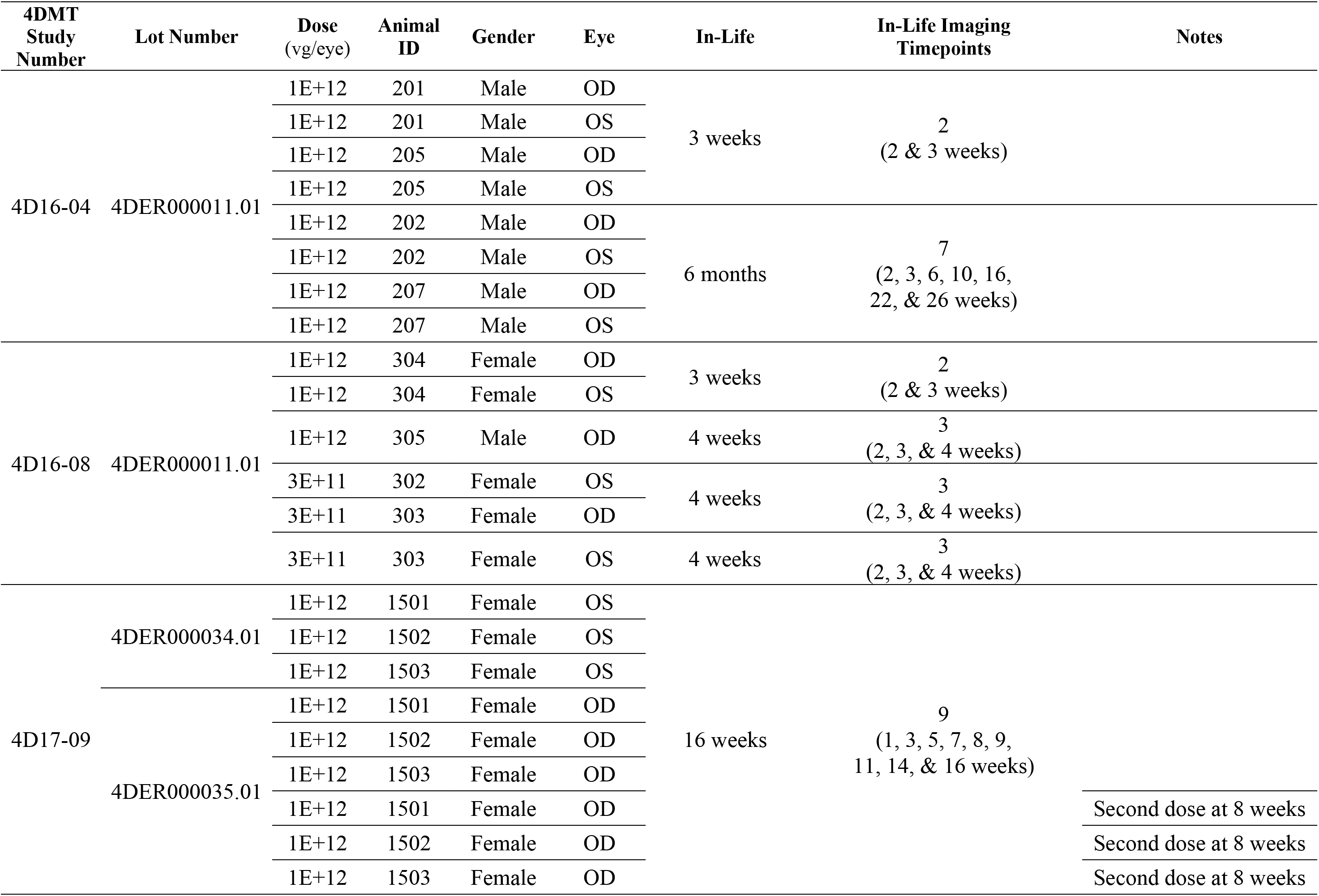
R100.CAG-EGFP Vector Characterization Studies

**Supplemental Table 3.** R100.CAG-anti-VEGF scFv Choroidal Neovascularization Study

| 4DMT<br>Study<br>Number | Lot Number | Animal<br>ID | Gender | Eye(s) | Dose | In-Life Imaging<br>Timepoints | Notes |
| --- | --- | --- | --- | --- | --- | --- | --- |
| 4D19-15 | N/A | B419 | Male | OU | vehicle | 2<br>(2 & 4 weeks post-laser) |  |
|  |  | B170 | Male | OU |  |  |  |
|  |  | B165 | Female | OU |  |  |  |
|  |  | B055 | Female | OU |  |  |  |
|  |  | A729 | Female | OU |  |  |  |
|  |  | A781 | Female | OU |  |  |  |
|  | 4DEP000013.01 | B412 | Male | OU | 1E+11 vg/eye |  |  |
|  |  | B416 | Male | OU |  |  |  |
|  |  | A752 | Female | OU |  |  |  |
|  |  | B142 | Female | OU |  |  |  |
|  |  | B050 | Female | OU |  |  |  |
|  |  | A700 | Female | OU |  |  |  |
|  | 4DEP000013.01 | B216 | Male | OU | 1E+12 vg/eye |  |  |
|  |  | B418 | Male | OU |  |  |  |
|  |  | B085 | Female | OU |  |  |  |
|  |  | A645 | Female | OU |  |  |  |
|  |  | A979 | Female | OU |  |  |  |
|  |  | B267 | Male | OU |  |  |  |

