## Supplementary figures and images for "Directed Evolution of AAV Targeting Primate Retina by Intravitreal Injection Identifies R100, a Variant Demonstrating Robust Gene Delivery and Therapeutic Efficacy in Non-Human Primates"

### Supplemental Figure S1

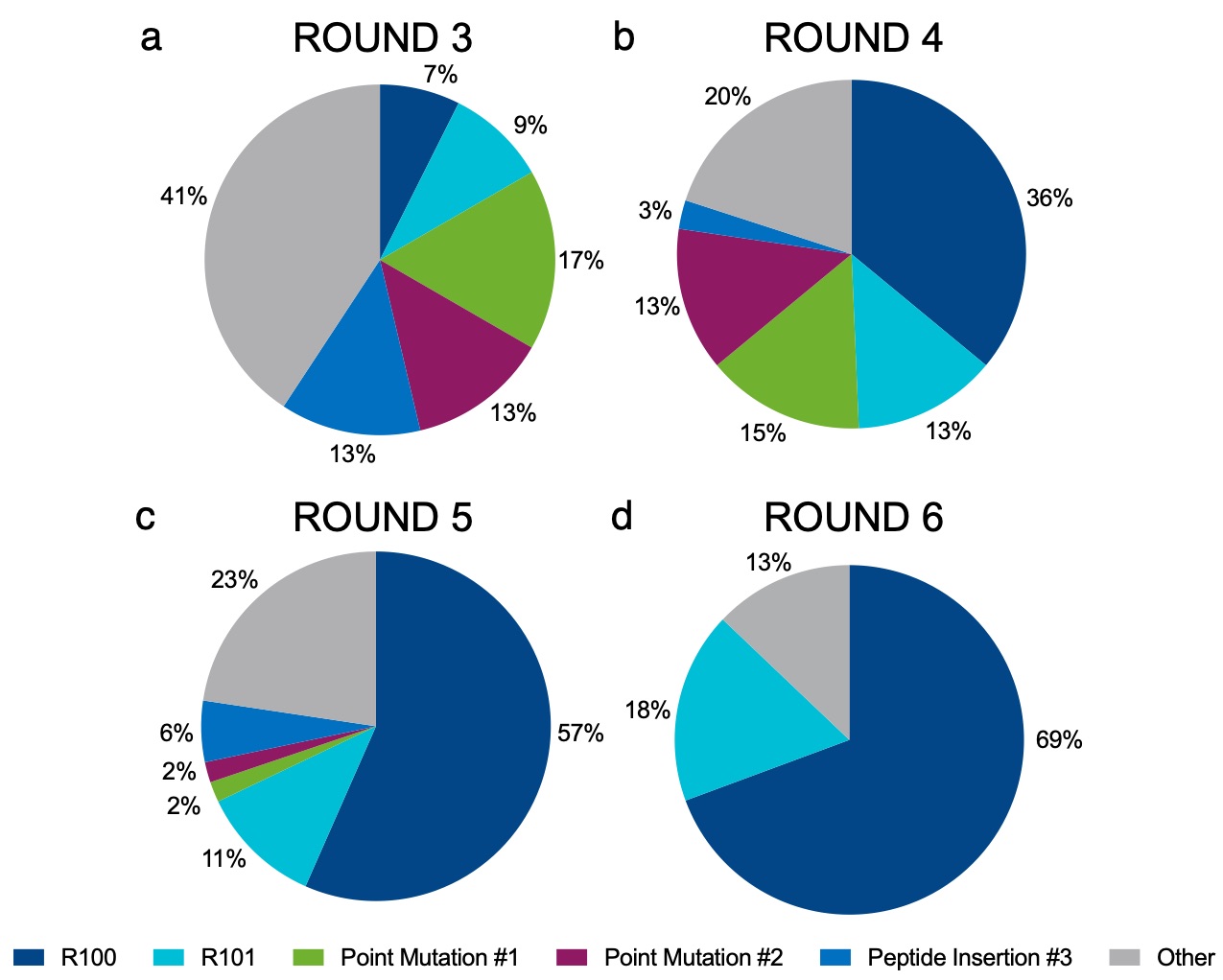

### Supplemental Figure S2

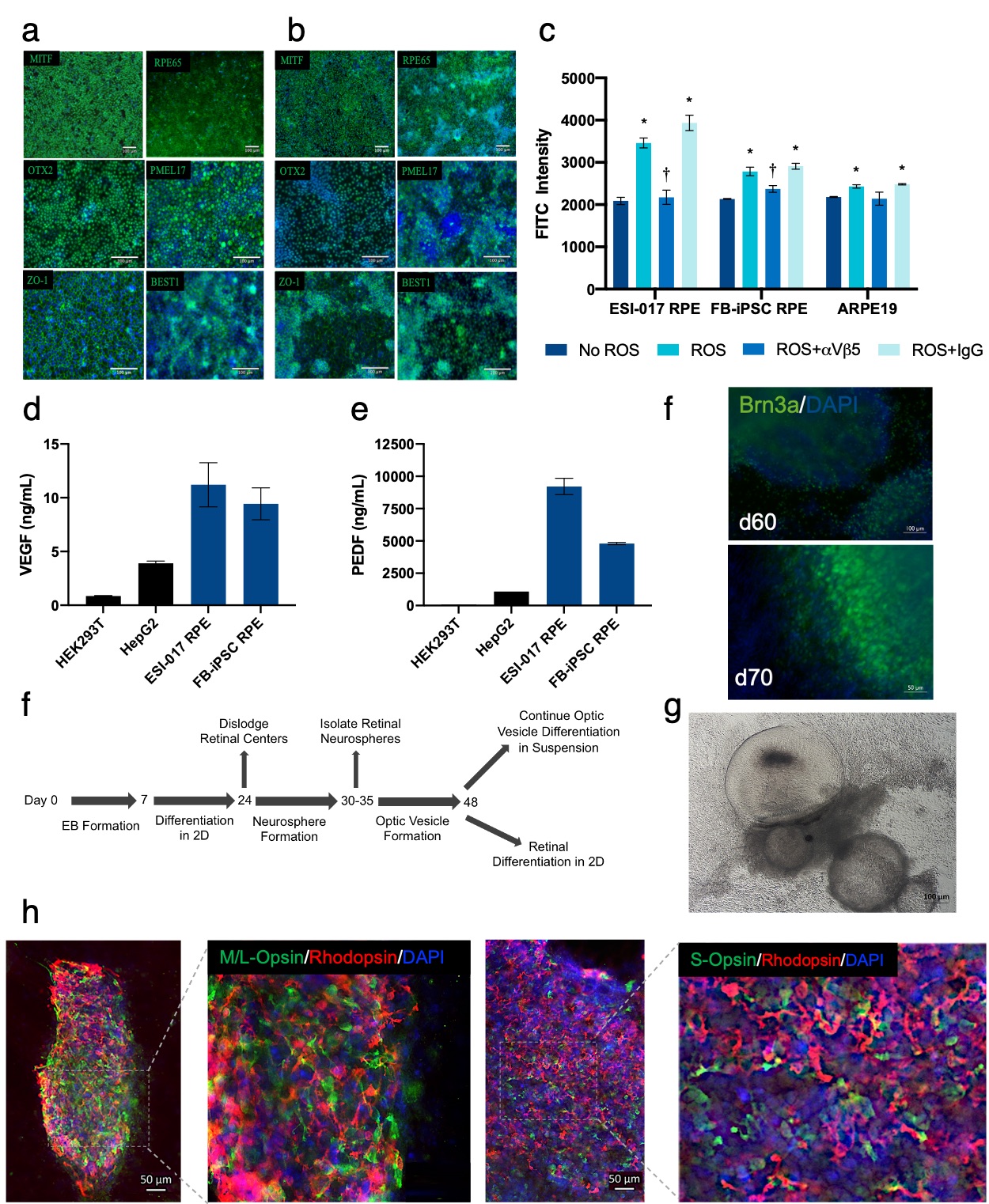

### Supplemental Figure S3A

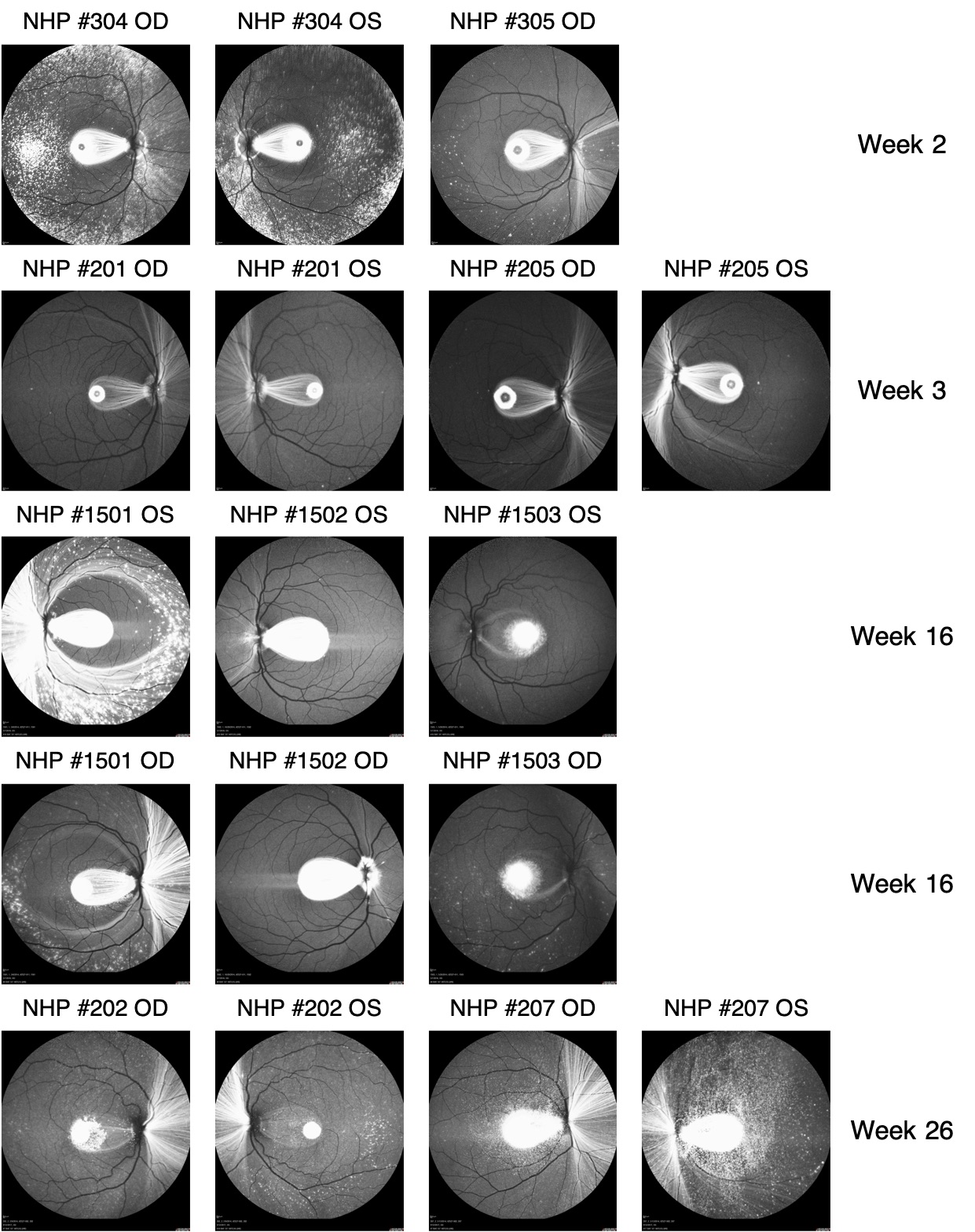

### Supplemental Figure S3B

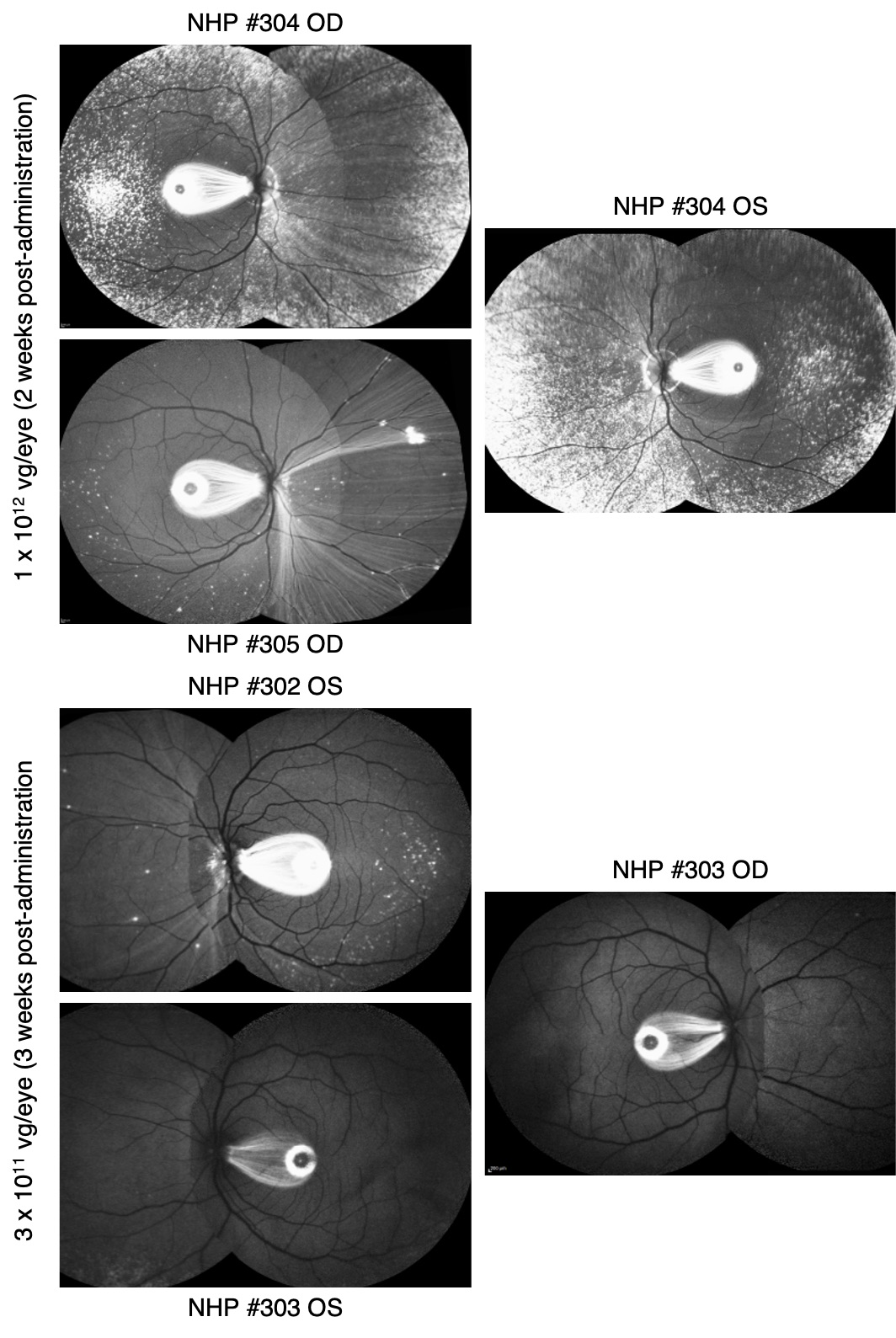

### Supplemental Figure S4

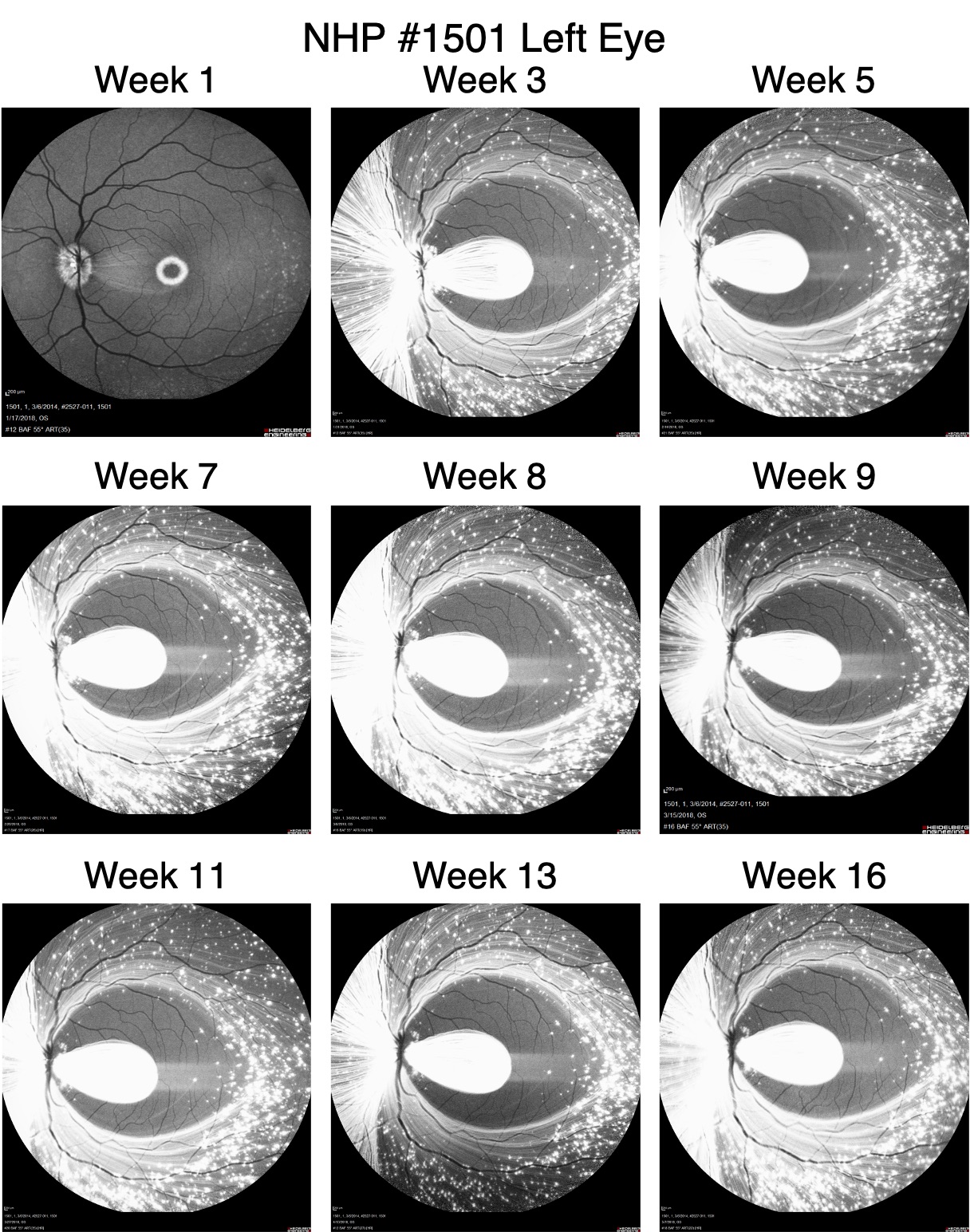

### Supplemental Figure S5A

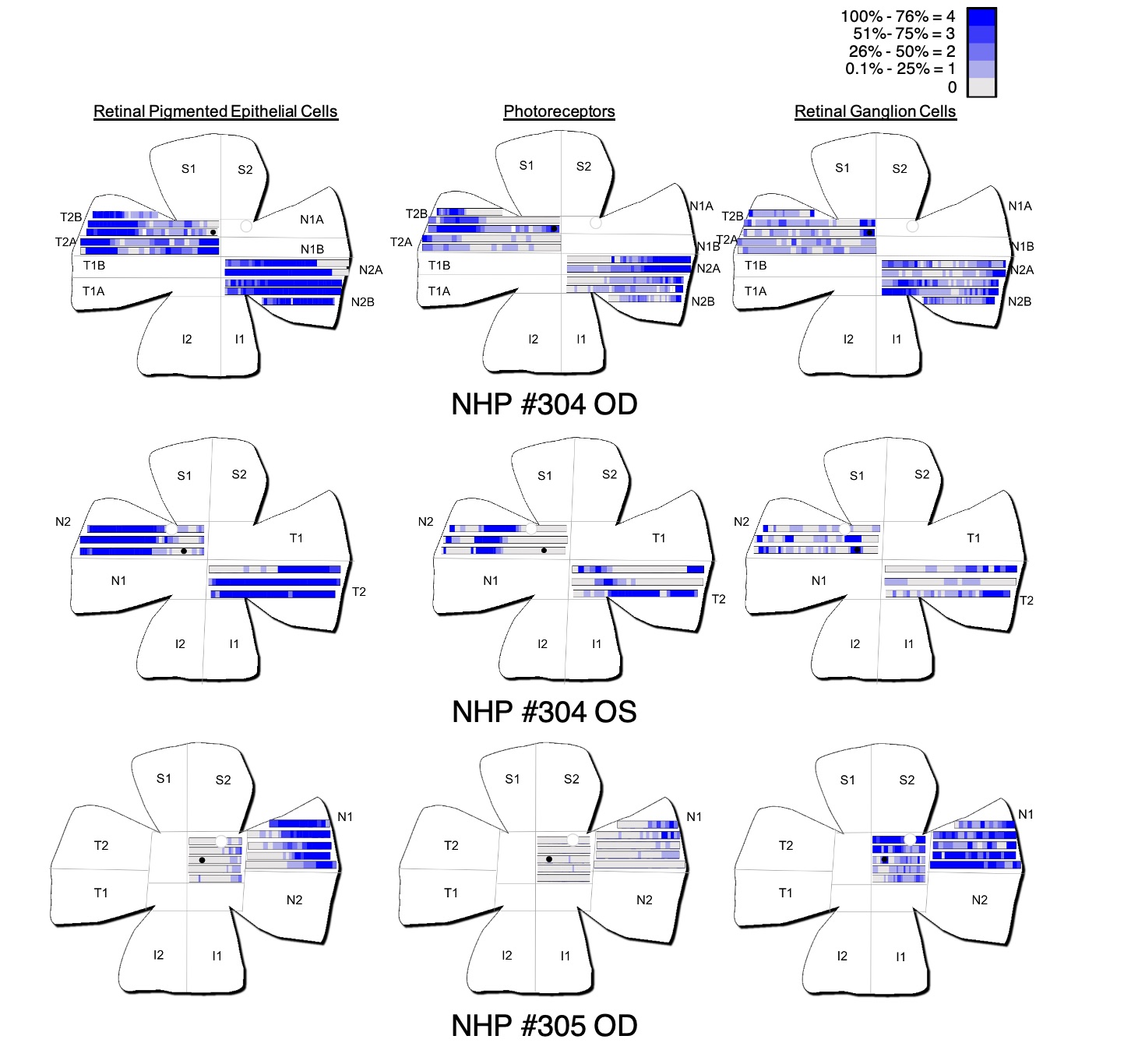

### Supplemental Figure S5B

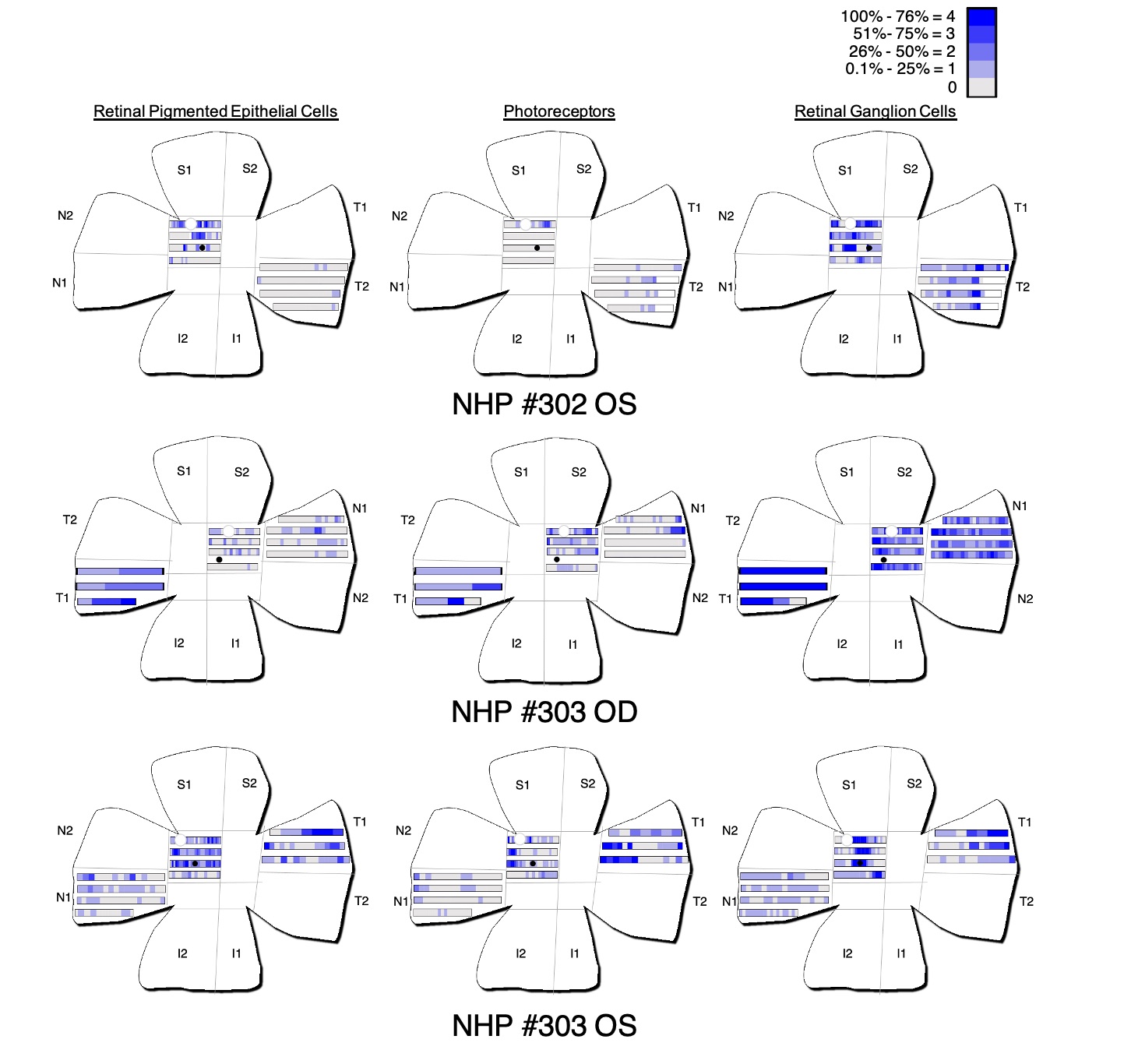

### Supplemental Figure S6

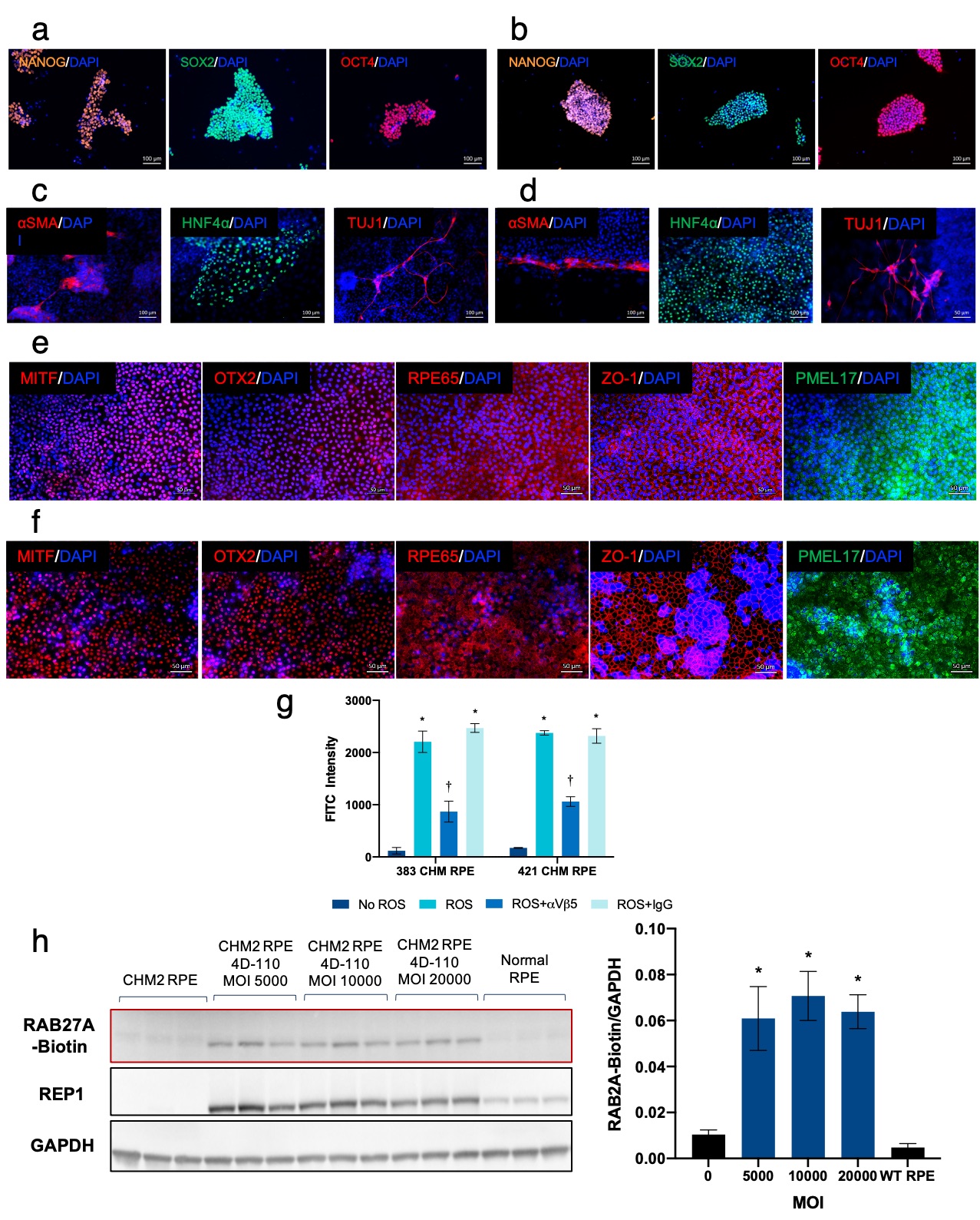

### Supplemental Figure S7

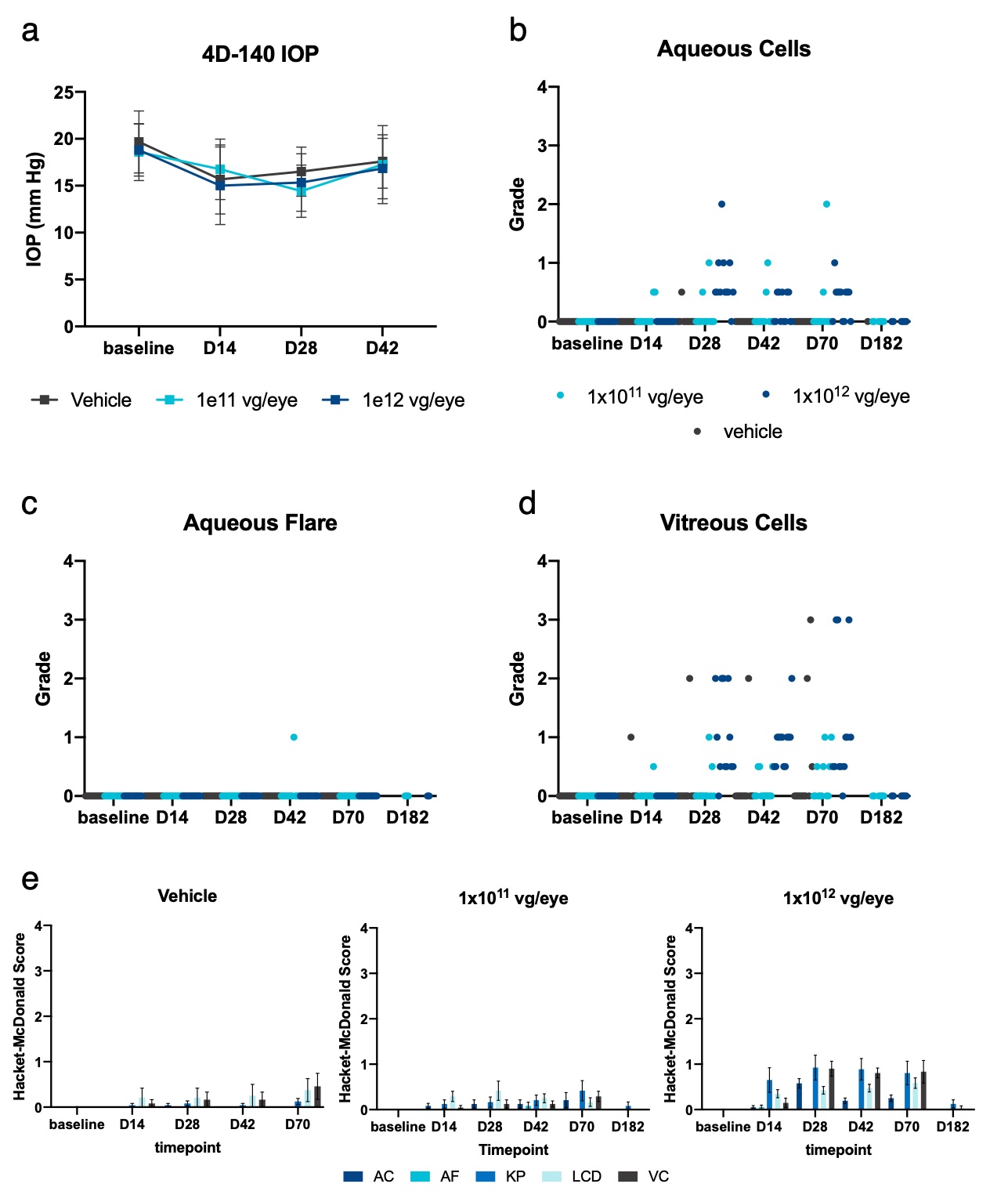

### Supplemental Figure S8

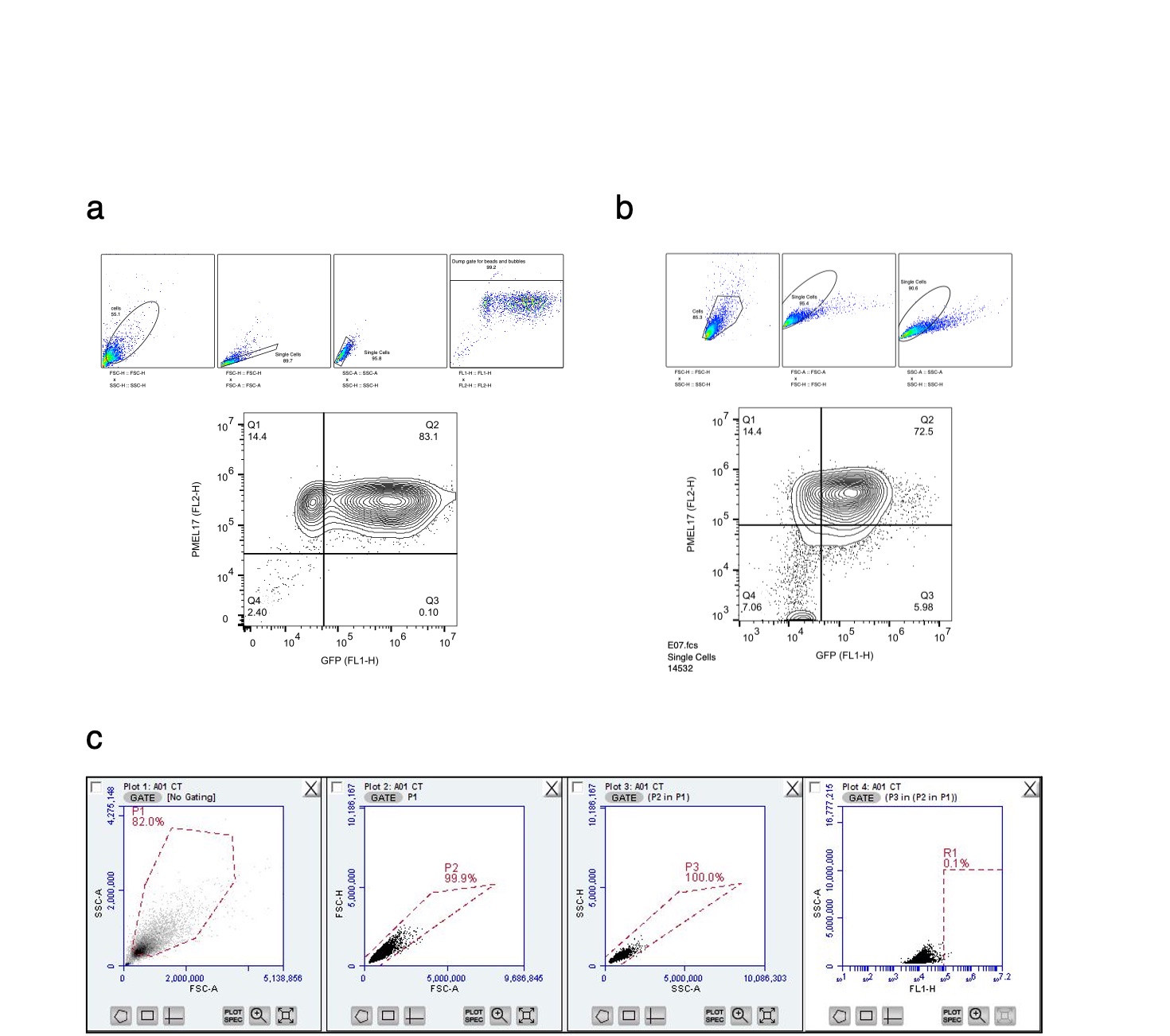

### Supplemental Figure S9

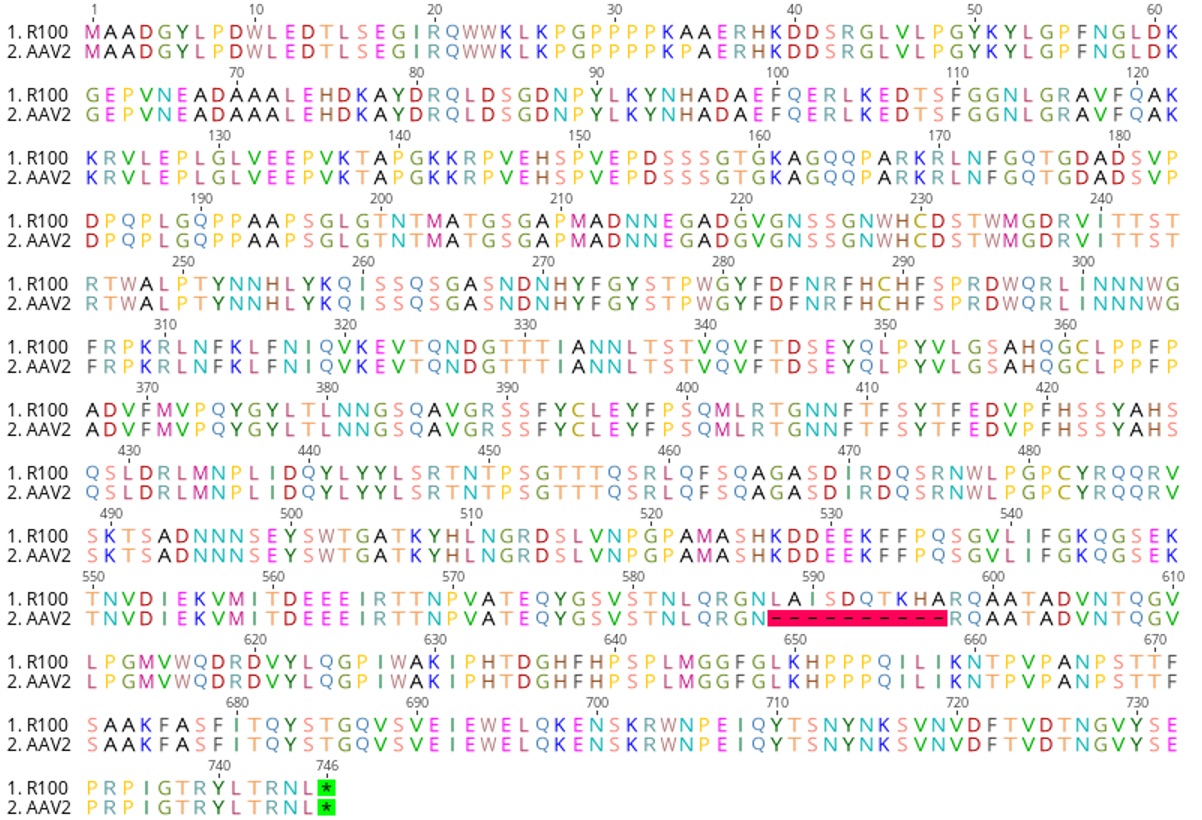
